# Redesigning the *Drosophila* histone gene cluster: An improved genetic platform for spatiotemporal manipulation of histone function

**DOI:** 10.1101/2024.04.25.591202

**Authors:** Aaron T. Crain, Markus Nevil, Mary P. Leatham-Jensen, Katherine B. Reeves, A. Gregory Matera, Daniel J. McKay, Robert J. Duronio

**Affiliations:** Curriculum in Genetics and Molecular Biology, University of North Carolina, Chapel Hill, NC, 27599 USA; Integrative Program for Biological and Genome Sciences, University of North Carolina, Chapel Hill, NC, 27599 USA; Seeding Postdoctoral Innovators in Research & Education, University of North Carolina, Chapel Hill, NC 27599 USA; Department of Biology, University of North Carolina, Chapel Hill, NC, 27599 USA; Department of Genetics, University of North Carolina, Chapel Hill, NC, 27599 USA; Lineberger Comprehensive Cancer Center, University of North Carolina, Chapel Hill, NC, 27599 USA

## Abstract

Mutating replication-dependent (RD) histone genes is an important tool for understanding chromatin-based epigenetic regulation. Deploying this tool in metazoan models is particularly challenging because RD histones in these organisms are typically encoded by many genes, often located at multiple loci. Such RD histone gene arrangements make the ability to generate homogenous histone mutant genotypes by site-specific gene editing quite difficult. *Drosophila melanogaster* provides a solution to this problem because the RD histone genes are organized into a single large tandem array that can be deleted and replaced with transgenes containing mutant histone genes. In the last ∼15 years several different RD histone gene replacement platforms have been developed using this simple strategy. However, each platform contains weaknesses that preclude full use of the powerful developmental genetic capabilities available to *Drosophila* researchers. Here we describe the development of a newly engineered platform that rectifies many of these weaknesses. We used CRISPR to precisely delete the RD histone gene array (*HisC*), replacing it with a multifunctional cassette that permits site-specific insertion of either one or two synthetic gene arrays using selectable markers. We designed this cassette with the ability to selectively delete each of the integrated gene arrays in specific tissues using site-specific recombinases. We also present a method for rapidly synthesizing histone gene arrays of any genotype using Golden Gate cloning technologies. These improvements facilitate generation of histone mutant cells in various tissues at different stages of *Drosophila* development and provide an opportunity to apply forward genetic strategies to interrogate chromatin structure and gene regulation.

## Introduction

The organization and regulation of genetic information in eukaryotes is coordinated by the packaging of DNA and core histone proteins (H2A, H2B, H3, H4) into nucleosomes to form chromatin. Nucleosomes are thought to serve as regulatory hubs for signals that direct chromatin folding and gene expression via post-translational modification (PTM) of the highly conserved histone N-terminal tail residues (Strahl and Allis 2000; Kouzarides 2007). A complex pattern of histone PTMs is installed, removed, and interpreted by *trans*-acting factors that “write”, “erase”, and “read” histone PTMs. The biological functions of histone PTMs are usually inferred by manipulating the activity of readers, writers, and erasers. However, chromatin proteins often have multiple targets in addition to histones, which complicates the determination of histone PTM function using this strategy. For example, the Brahma subunit of BAP and PBAP plays a dual role by reading H3 Lysine 14 acetylation (H3K14ac) and enhancing a writer of H3K27 acetylation, thereby contributing to many downstream effects (Kimura *et al*. 2003; Tie *et al*. 2012). Likewise, the H4K20 methyltransferase Set8 has non-histone targets in mammals (Shi *et al*. 2007; Weirich *et al*. 2015). Last, the TIP60 complex acetylates up to five lysines on the same Histone H4 tail, masking the contribution of any single acetylated H4 lysine via mutation of TIP60 (Jacquet *et al*. 2016). When manipulating proteins that act on chromatin, we must remain aware of pleiotropic phenotypes that can obfuscate the true function of individual histone PTMs. Consequently, we and others have turned to directly mutating histone tail residues in *Drosophila melanogaster* to understand histone PTM function more definitively (Günesdogan *et al*. 2010; McKay *et al*. 2015; Zhang *et al*. 2019).

*Drosophila melanogaster* is an excellent experimental organism for manipulating the chromatin environment through direct mutation of histone genes (Corcoran and Jacob 2023). The organization of replication-dependent (RD) histone genes in *Drosophila* makes histone genetics tractable. Whereas in most well-studied metazoans (e.g. *Homo sapiens, Danio rerio, Xenopus laevis*, and *Caenorhabditis elegans*) the RD histone genes are located in clusters distributed at multiple loci (Figure 1A), *D. melanogaster* carry a single RD histone locus on chromosome 2L (*HisC*) consisting of a 5 kb histone gene unit that is tandemly arrayed about 100 times (Figure 1B) (Lifton *et al*. 1978; McKay *et al*. 2015; Bongartz and Schloissnig 2019). Several groups have taken advantage of this genomic feature to engineer histone genotypes in *Drosophila* by combining a homozygous *HisC* deletion with transgenic RD histone gene arrays encoding mutant histone proteins (Figure 2). Importantly, observations made in *D. melanogaster* are readily transferable to other organisms because the core histones are highly conserved among metazoans. The average amino acid identity for the core histone tails of H2a, H2b, H3, and H4 among five commonly studied metazoan species are 88.5%, 85.7%, 99.3%, and 99.5%, respectively (Figure 1C).

**Figure 1.**
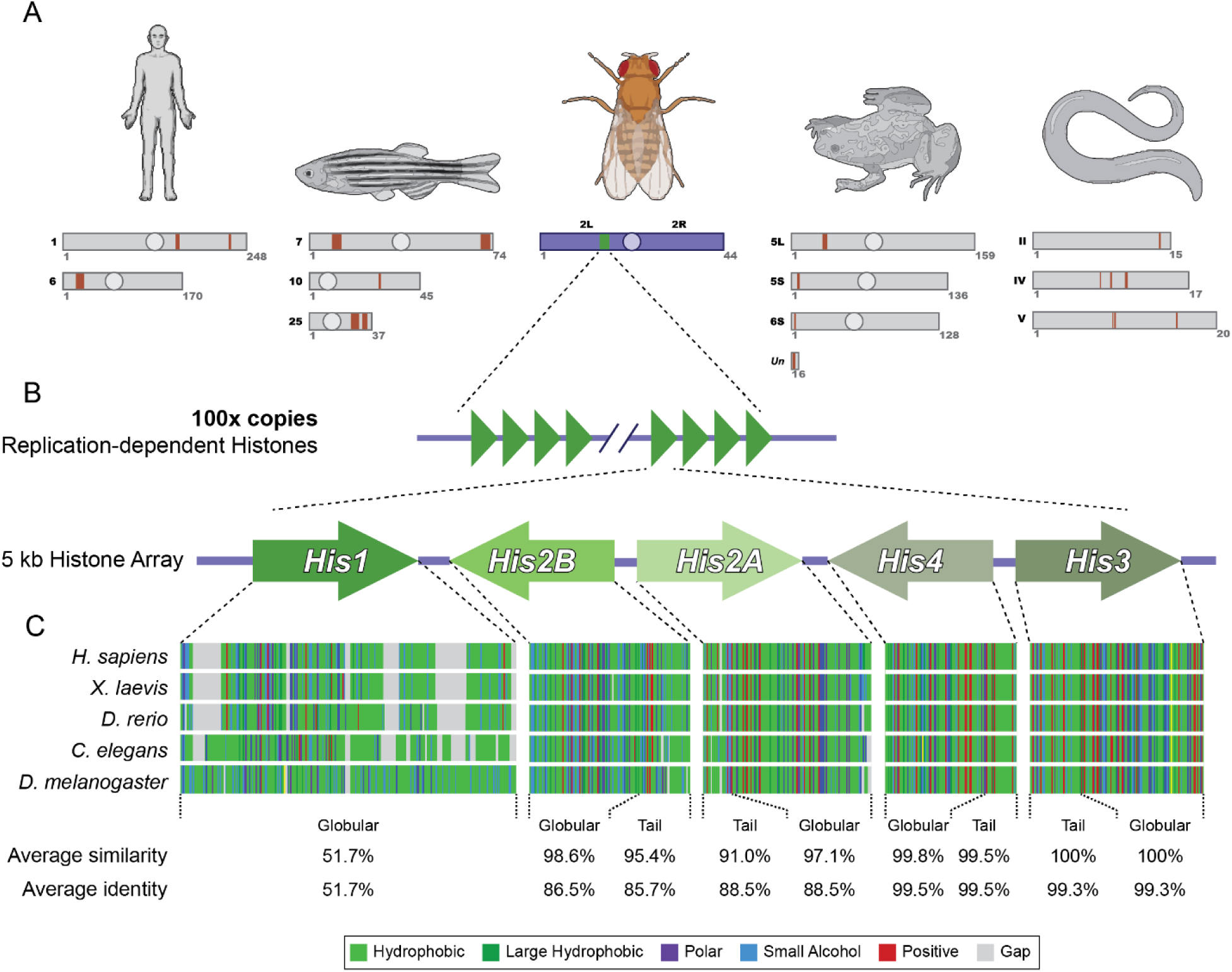
Comparison of replication-dependent histone genes and proteins among metazoan model organisms. A) Diagram depicting histone gene clusters in *H. sapiens*, *D. rerio*, *D. melanogaster*, *X. laevis*, and *C. elegans*. Only chromosomes containing histone gene clusters are shown, and histone gene clusters are represented by orange boxes for all organisms except *Drosophila*, where the histone gene cluster is represented by a green box. B) *D. melanogaster* possesses ∼100 tandem-repeats of a nearly identical, 5kb histone gene unit containing each of the five replication-dependent histone genes: *H1*, *H2b*, *H2a*, *H4*, and *H3*. C) Multiple sequence protein alignment of the five replication-dependent histones from each of the listed organisms. Amino acids are colored by identity (Hydrophobic = light green, Large Hydrophobic = dark green, Polar = purple, Small Alcohol = blue, Positive = red) (Brown et al., 1998) and gaps are represented as grey boxes. Average amino acid similarity and identity were calculated for each species compared to *H. sapiens*.

**Figure 2.**
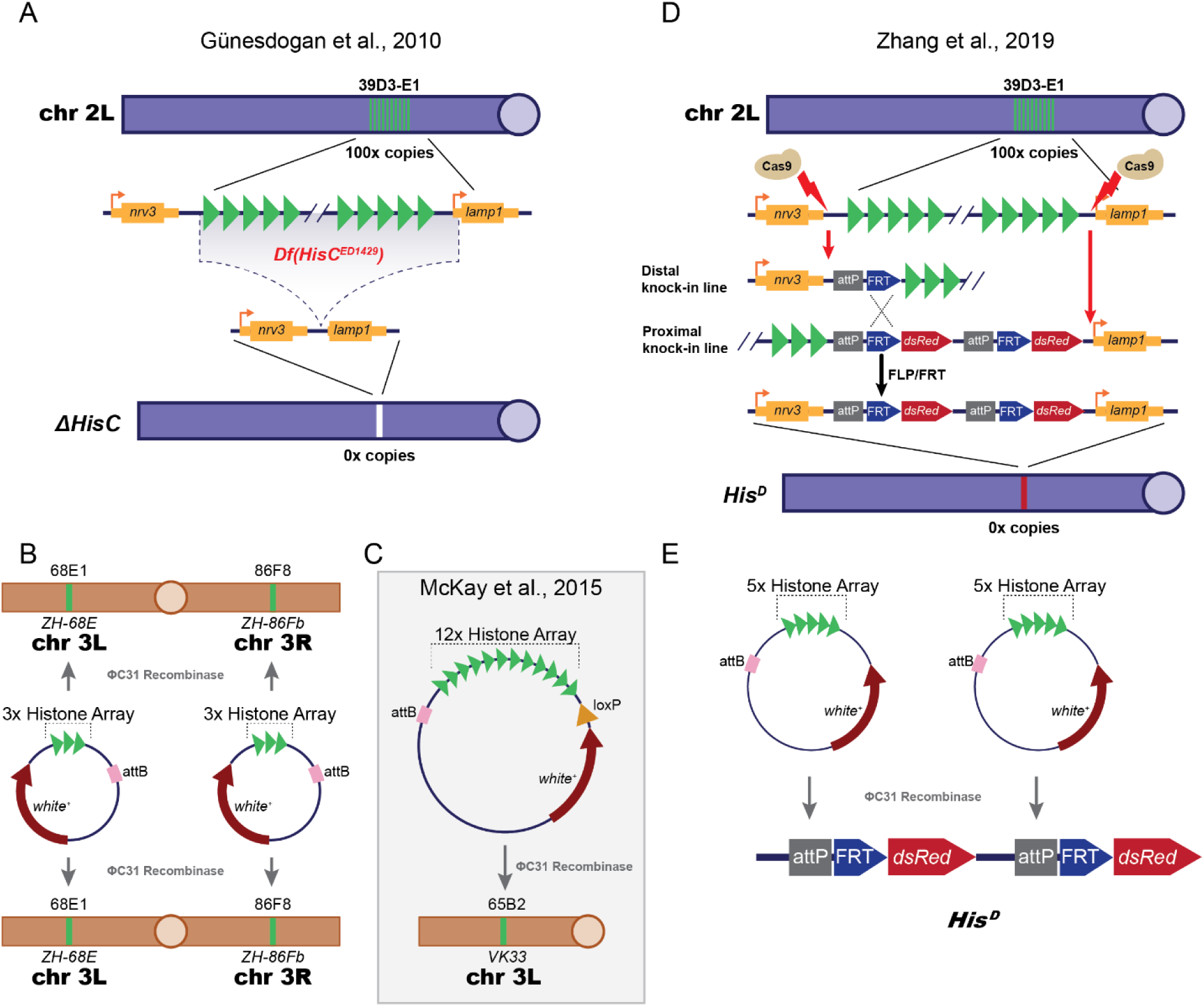
Summary of existing histone replacement strategies in *Drosophila melanogaster*. A) Diagram depicting the generation of *Df(HisC^ED1429^)* (*ΔHisC*) by Günesdogan et al., 2010. Wild-type chromosome 2L (top) is shown in purple with the centromere shown in light purple. The replication-dependent histone cluster is shown as vertical green lines. A zoom-in of the RD histone cluster (middle) depicts the region of chromosome 2L that is deleted in *ΔHisC* (dotted lines filled with grey), which includes the upstream regulatory region, transcription start site, and 26 bp of the 5’ UTR of the *lamp1* gene. Depiction of chromosome 2L after removal of the replication-dependent histone cluster in *ΔHisC* (bottom). B) Diagram depicting histone replacement strategy developed by Günesdogan et al., 2010. Chromosome 3 is shown in orange with the centromere shown in light orange. ΦC31-mediated integration sites (ZH-68E and ZH-86Fb) are shown as green bars at their approximate locations on chromosome 3 (68E1 and 86F8) (Bischof *et al*. 2007). Four different 3x histone gene unit transgenes were used for rescue of homozygous *ΔHisC.* C) Diagram depicting the single-site histone replacement strategy developed by McKay et al., 2015. Chromosome 3L is shown in orange with the centromere shown in light orange. The ΦC31-mediated integration site (VK33) is shown as a green bar at the approximate location on chromosome 3 (65B2). D) Diagram depicting the generation of the *His^D^* deletion by Zhang et al. 2019 by independent CRISPR-Cas9 knock-ins flanking the endogenous histone array. *His^D^* was obtained by FLP/FRT recombination between the two CRISPR engineered chromosomes to remove the endogenous histone locus. E) Diagram depicting the histone replacement strategy developed by Zhang et al. 2019, where two 5x Histone array BACs are integrated at the *His^D^* locus.

Although considerable effort went into developing histone gene replacement platforms in *Drosophila*, these technologies have limitations. Herzig and colleagues developed the first RD histone replacement platform by deleting the entire *HisC* locus using the DrosDel system (Ryder et al. 2004), thereby generating *Df(HisC^ED1429^)* which we refer to hereafter as “*ΔHisC*” (Figure 2A) (Günesdogan *et al*. 2010). To rescue the lethal phenotype of a *ΔHisC* homozygote, four plasmid-based transgenes bearing three tandemly repeated 5kb histone gene units containing each of the five RD histone genes (i.e. a 3x array of the *H1*, *H2a*, *H2b*, *H3*, and *H4* genes) were inserted into the genome at two different loci using ΦC31 Recombinase (Figure 2B). Interestingly, the authors found that a total of 12 copies of transgenic, wild-type histone gene units (∼6% of the endogenous number) were sufficient to support viability and fertility of homozygous *ΔHisC* animals. This platform allowed for the first direct interrogation of histone residue function in metazoan development (Hödl and Basler 2012; Pengelly *et al*. 2013). Unfortunately, this method was ultimately limited in ease-of-use because of the need to generate and manipulate transgenes at four different loci. Thus, a further development by McKay et al. 2015 generated a BAC-based platform that could rescue deletion of the endogenous histone locus with a single 12x histone gene array inserted at one locus (Figure 2C). These improvements simplified the construction of transgenic genotypes and downstream genetics. Subsequently, a histone replacement platform was developed using CRISPR-Cas9-mediated engineering of the *HisC* locus (Figure 2D), replacing the endogenous histone gene array with two transgenic 5x histone gene arrays (Zhang et al., 2019; Figure 2E). These three gene replacement platforms have been used to greatly expand our understanding of metazoan histone residue function in the context of animal development (Günesdogan *et al*. 2010; Hödl and Basler 2012; Pengelly *et al*. 2013, 2015; McKay *et al*. 2015; Yung *et al*. 2015; Penke *et al*. 2016; Graves *et al*. 2016; Meers *et al*. 2017, 2018a; b; Armstrong *et al*. 2018, 2019; Copur *et al*. 2018; Leatham-Jensen *et al*. 2019; Zhang *et al*. 2019; Koreski *et al*. 2020; Finogenova *et al*. 2020; Regadas *et al*. 2021; Lindehell *et al*. 2021; Crain *et al*. 2022; Corcoran and Jacob 2023; Salzler *et al*. 2023).

Each of these RD histone gene replacement strategies have disadvantages that preclude making full use of the arsenal of genetic tools available in *Drosophila melanogaster*. Therefore, we engineered a novel, designer *HisC* locus that combines useful features of existing histone replacement platforms with new capabilities, while maintaining a degree of backwards compatibility to complement previous work. We have added new functionalities to *HisC* that expand utility and afford flexibility in histone genotype engineering. Together this new system provides a more sophisticated toolkit for generating complicated genetic backgrounds, carrying out forward genetics, and performing studies requiring spatiotemporal control of mutant histone gene expression. Here, we describe the features and demonstrate intended uses of this new system.

## Results

### Engineering a multi-functional replication-dependent histone gene deletion in

#### Drosophila melanogaster

The endogenous chromatin context has a direct effect on the expression of genes, and thus may impact the expression of RD histone genes inserted at ectopic genomic locations. Therefore, when designing our new platform, we targeted the *HisC* locus for CRISPR-Cas9 genome editing such that inserted histone genes would be expressed in their endogenous context. We used *ΔHisC* as our substrate allele for CRISPR engineering to avoid the need to repair across a 0.5 megabase region (i.e. on either side of the endogenous 100x histone gene array). We designed a CRISPR-Cas9 guide RNA targeting genomic sequence near the centromere distal 3’ P-element sequence located at *ΔHisC*, and another guide RNA targeting the 5’ UTR of the centromere proximal *lamp1* gene (Figure 3A). We constructed a repair template containing *Actin5C-dsRed* between distal and proximal homology arms to facilitate repair of the 5.6 kilobase gap resulting from site-specific cleavage by Cas9 (Figure 3B). Potential recombinants were obtained by recovering dsRed-positive progeny from G0 animals containing a transformed germ line. Sequencing of *HisC* PCR products isolated from transformants confirmed a clean repair that extended beyond the homology arms of the repair template into the adjacent, exogenous sequence (Figure 3C). We also confirmed the intended mutation of the guide RNA sites to abrogate re-cutting of repaired loci (Figure 3C). A PCR product spanning the locus was analyzed by Oxford Nanopore long read sequencing and confirmed the presence of the entire repair template (Figure 3D). As this newly engineered locus is designed with many desirable features, we named it the *ΔHisC^cadillac^* locus. As expected, *ΔHisC^cadillac^* is inviable when homozygous and a 12x histone wild-type (HWT) transgene located on the third chromosome rescues this lethality, demonstrating the applicability of *ΔHisC^cadillac^* for a new histone gene replacement platform.

**Figure 3.**
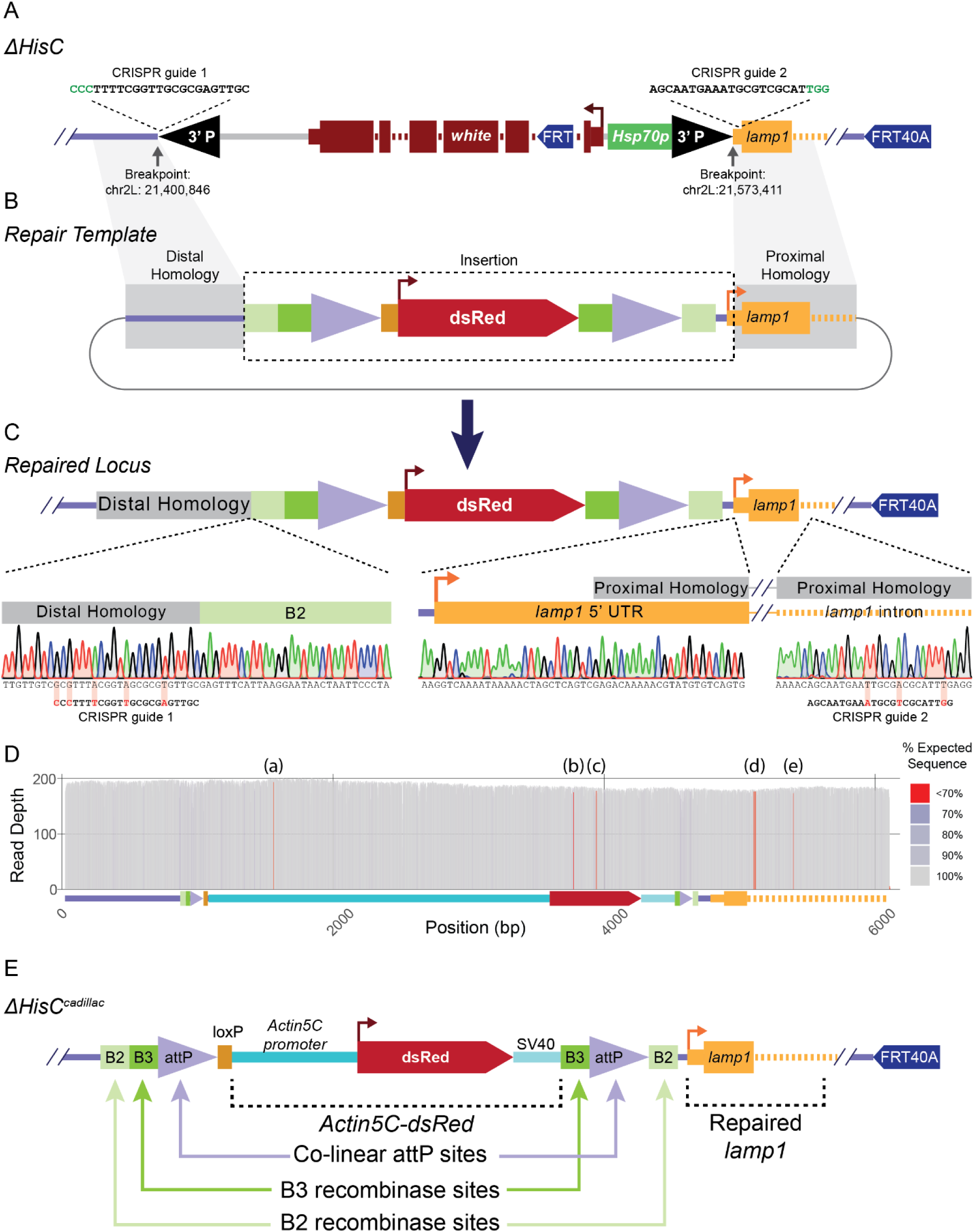
Engineering of *ΔHisC^cadillac^*, a novel designer histone deletion locus in *Drosophila melanogaster* as a platform for histone gene replacement. A) Diagram of the *ΔHisC* locus with locations of the CRISPR-Cas9 guideRNA target sequences used for gap repair. *ΔHisC* is marked by *mini-white* (*white*) with an FRT site integrated at the first intron. Breakpoints between endogenous sequence and P-element sequence are noted using dm6 coordinates (Hoskins *et al*. 2015). The centromere-proximal breakpoint results in removal of the transcription start site and upstream regulatory region plus 26 bp of the 5’UTR of *lamp1*. B) A simplified diagram of the CRISPR-Cas9 repair template plasmid, noting the distal and proximal homologies and desired insertion sequence. C) A simplified diagram of the repaired locus with Sanger sequence traces over the mutated guideRNA sequences and junctions between endogenous sequence (distal/proximal homologies) and inserted sequences (B2, and 5’-UTR of *lamp1*). D) Oxford Nanopore long read sequence validation of the *ΔHisC^cadillac^* locus. Read depth at each position of the locus is represented on the y-axis with the color indicating the percent of reads with the expected sequence. The red positions (a), (b), (c), (d), and (e) represent differences between *ΔHisC^cadillac^* and the reference sequence. E) Fully annotated diagram of the *ΔHisC^cadillac^* locus.

*ΔHisC^cadillac^* contains several features that greatly expand the genetic toolkit at the *Drosophila* RD histone locus (Figure 3E). First, it precisely deletes *HisC*, leaving the *lamp1* gene intact, in contrast to the widely used *ΔHisC* allele which deletes the *lamp1* promoter. Although *lamp1* is not required for viability, it encodes a protein critical for lipid metabolism (Chaudhry *et al*. 2022). Therefore, *ΔHisC^cadillac^* eliminates any potential confounding phenotypes resulting from loss of *lamp1*. *ΔHisC^cadillac^* contains two co-linear *attP* sites for integration of either one or two transgenic histone gene arrays. This design also allows for combinatorial integration of two arrays with different genotypes and integration via recombinase-mediated cassette exchange (RMCE). Inclusion of *Actin5C-dsRed* simplifies scoring of genotypes through positive identification of the *ΔHisC^cadillac^* chromosome (Supplemental Figure 1A). The *attP* sequences and *Actin5C-dsRed* are flanked by B2 and B3 recombination sites, which allow for excision of sequences integrated at the *attP* sites using UAS/Gal4-driven B2 and B3 site-specific recombinases (Nern *et al*. 2011). Importantly, the location of these B2 and B3 recombination sites allows for the removal of i) only sequences integrated into the distal *attP* site via B3R or ii) sequences integrated into either or both of the *attP* sites via B2R. To expand the combinatorial capacity of the locus a *loxP* site was inserted between the *attP* sites. This *loxP* site can be used to remove vector sequences after integration (McKay *et al*. 2015; Meers *et al*. 2018a). Last, the *ΔHisC^cadillac^* chromosome also contains the centromere-linked FRT40A to facilitate mitotic recombination. The following sections describe the utility of these and other features of *ΔHisC^cadillac^*.

### Generating histone mutant mosaics in the adult *Drosophila* eye for use in forward genetic screens

A primary draw of histone gene replacement strategies is the ability to assess phenotypes of completely mutant animals. However, due to the importance of many histone residues, finely tuned assessment of effects in certain tissues (e.g. those in adults or pupae) is sometimes precluded by the failure of mutant animals to develop to the necessary stage. Generating genetically mosaic tissues in flies that otherwise remain heterozygous for a lethal histone mutation provides a solution to this problem. We therefore created a strategy for generating mosaic eye tissue composed of clones of cells with different histone genotypes. The *Drosophila* eye has an easily observable color and structural organization; it is dispensable for survival and can be manipulated using a variety of genetic tools (St Johnston 2002; Baker *et al*. 2014). Inducing mitotic recombination during development results in a mosaic tissue with clones of homozygous wild-type and clones of homozygous mutant cells that directly compete against one another as the eye grows. For instance, Kanda and colleagues showed that eye clones mutant for the histone methyltransferase *trithorax related* have a growth advantage over their wild-type neighbors (Kanda *et al*. 2013). However, such experiments are difficult to perform for histone mutant cells using the existing histone gene replacement platforms.

The primary complication is the *white* (*w*) marker gene within *ΔHisC* (Günesdogan *et al*. 2010). In most experiments, mutant adult eye clones are also *w* mutant (and thus white in color) and wild-type eye clones are *w*+ (and thus red in color). Because the *ΔHisC* allele carries a *white* marker gene (Günesdogan *et al*. 2010; McKay *et al*. 2015), *ΔHisC* homozygous cells resulting from mitotic recombination are *w+*, obscuring the identification of histone mutant clones, especially when they are small. That is, none of the resulting histone mutant clones lack *white* expression. Furthermore, we found that flies expressing *ey-FLP* in a background with *ΔHisC* but no other FRT-containing chromosome had red-and-white mosaic eyes, likely due to intrachromosomal excision of sequences between the FRT sites located in *ΔHisC* and the FRT sites located at 40A (data not shown; FRT sites diagramed in Figure 3A).

We engineered *ΔHisC^cadillac^* to solve these problems and to allow easy and efficient assessment of clones resulting from mitotic recombination, particularly in the eye. The *white* gene of *ΔHisC* is replaced by *Actin5C-dsRed* in *ΔHisC^cadillac^*, providing a bright marker of the locus under fluorescent light, yet imparting no discernable eye color under white light (Supplemental Figure 1A). To test whether *ΔHisC^cadillac^* could be used to detect histone mutant mosaic tissue using fluorescent marker genes rather than *white*, we first created two constructs containing either *Act5C-sfGFP* or *3xP3-sfGFP* and lacking *white*. We independently inserted these constructs into *attP40* on chromosome 2L, followed by recombination to a chromosome containing FRT40A (Supplemental Figure 1B). Neither *sfGFP*-containing transgene has discernable eye color under white light but are distinguishable with a fluorescent microscope, where *Act5C-sfGFP* is expressed in the entire animal and *3xP3-sfGFP* is expressed only in the eyes (Supplemental Figure 1C).

With these new chromosomes, we used *ey*-FLP to drive mitotic recombination at FRT40A, thereby generating sister clones in the adult that are either genotypically *ΔHisC^cadillac^*/*ΔHisC^cadillac^* or *HisC*^+^/*HisC*^+^, both of which contain a 12x histone gene array (12x HWT) on chromosome 3 (Figure 4A). In this way we can assess the proliferation of cells that contain only 12x HWT (dsRed+, sfGFP-), cells that contain 212 histone gene copies (dsRed-, sfGFP+) or cells with 112 histone gene copies (dsRed+, sfGFP+) (Figure 4A). Homozygous *ΔHisC^cadillac^* cells fail to proliferate and do not contribute to the adult eye in the absence of a rescuing transgene (Figure 4B,C). This proliferation defect is rescued by the 12 copies of wild-type histones in the 12x HWT transgene (Figure 4B,C). Note that because *ey*-FLP is activated early in development and is expressed continuously in the eye, essentially the entire eye is populated by clones of cells with either 12x histone genes (magenta) or 212 histone genes (green).

**Figure 4.**
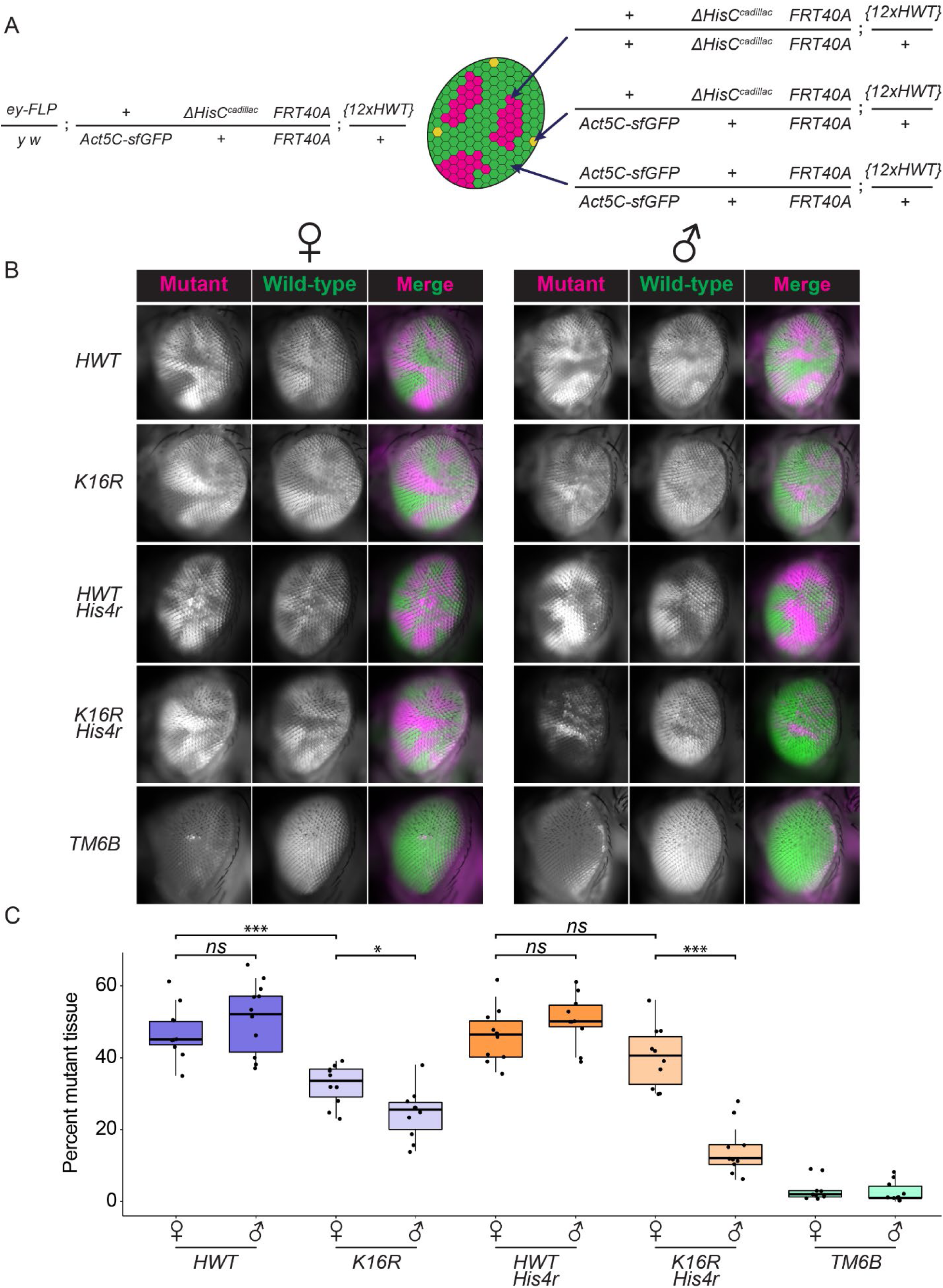
Fluorescent adult eye mosaic analysis of histone mutant genotypes. A) Left: genotype of fly in which mosaicism was induced using *ey-*FLP mediated mitotic recombination. Right: diagram of the expected clonal genotypes and fluorescent phenotypes after mitotic recombination during eye development. B) Female and male examples of mosaic adult *Drosophila* eyes containing *ΔHisC* homozygous clones rescued with the indicated 12x histone transgenes, with or without deletion of *His4r*. *TM6B* indicates flies lacking a 12x histone transgene. For each genotype the mosaic tissue is composed of clonal populations of cells that are homozygous for *Actin5C-sfGFP* (green; 200 endogenous histone genes and 12 transgenic), *ΔHisC^cadillac^* (magenta; 12 transgenic histone genes), or heterozygous (yellow; 100 endogenous histone genes and 12 transgenic). C) Quantification of the dsRed signal (homozygous *ΔHisC^cadillac^*) as a percentage of the total area of the eye image, displayed as box plots. Data separated into male (♂) and female (♀) flies. Blue boxes are genotypes with endogenous *His4r*, and orange boxes are genotypes without *His4r*. Green boxes indicate control tissue. Significance values calculated by Wilcoxon rank sum test: ns = not significant; * = p < 0.05; *** = p < 0.001.

Quantification of clone size indicates that 12x HWT clones grow equally well next to clones containing 212 copies of wild-type histones in both males and females (Figure 4C). This result was anticipated given that 12 copies of wild-type histones are sufficient for whole animal viability and fertility(Penke *et al*. 2016). Notably, the weak *white* expression from the HWT transgene rescue cassette does not obscure the visualization of dsRed or sfGFP in the adult eye.

The *Drosophila* compound eye is a powerful tool to study growth and cell proliferation through forward genetic screens using the FLP/FRT system (St Johnston 2002; Tapon *et al*. 2002; Pellock *et al*. 2007; Tseng *et al*. 2007; Kaufman 2017). We investigated whether we could measure proliferative differences in wild-type versus histone mutant cells using our fluorescent mosaic eye assay. This approach could unmask defects in the growth of histone mutant cells not apparent in whole animal mutants (Penke *et al*. 2016). During *Drosophila* imaginal disc development, cells that grow more slowly than their neighbors are outcompeted and actively killed (Morata and Ripoll 1975; Johnston 2009). Consequently, clones containing histone mutant cells that proliferate poorly will either be smaller than wild-type clones or absent from the adult eye altogether. We tested this premise using a previously described H4K16R substitution mutation (Armstrong et al., 2018). Because acetylated H4K16 is critical for upregulation of the X chromosome in XY males to balance gene expression with XX females, H4K16R mutant males are inviable whereas H4K16R females are viable (Armstrong et al., 2018). We asked to what extent the lethal phenotype of H4K16R mutant males might be due to defects in cell proliferation. We quantified the fraction of the adult eye composed of histone mutant tissue, with 50% representing no proliferation defect. We found that clones of H4K16R female tissue were slightly smaller than control (33.5% for H4K16R females compared to 45% for HWT females), and that this effect was more pronounced in males (25.5% for H4K16R males compared to 52% for HWT males) (Figure 4B, C). Interestingly, additional homozygous deletion of the single copy *His4r* gene, which is located outside of *HisC* and encodes an identical H4 protein, in H4K16R cells resulted in a more severe growth defect in male cells but did not affect female cells (12% for *H4^K16R^*, *His4r^null^* males versus 50% for *HWT, His4r^null^* males; 40.5% for *H4^K16R^, His4r^null^* females versus 46.5% for *HWT, His4r^null^* females) (Figure 4B,C). These data demonstrate that cells in the developing eye of males proliferate poorly when they can only produce H4K16R mutant histone. They also demonstrate that we can modulate the growth of histone mutant cells by mutating additional genes, indicating that this method could be utilized in forward genetic screens to identify new genes and pathways that regulate chromatin organization and epigenetic control of gene regulation and cell proliferation.

### PhiC31 integration of 4x, 6x, and 12x histone arrays into *ΔHisC^cadillac^*

The ability to insert transgenic histone gene arrays of various copy number and coding potential at the endogenous *HisC* locus is an essential feature of *ΔHisC^cadillac^*. We designed the locus to contain two co-linear *attP* sites for integration of *attB* containing constructs using established cloning and transgenic injection methods (Figure 5A) (McKay *et al*. 2015; Meers *et al*. 2018a). To test the integration efficacy at these sites, we took advantage of our “Designer Wild-type” (DWT) histone gene repeat, which contains polymorphisms in each of the 5 RD histone genes that allow us to distinguish endogenous wild-type histone genes from transgenic histone genes (Koreski *et al*. 2020). Using our previously described pMultiBAC vector (McKay *et al*. 2015), we obtained successful integration of 4x, 6x, and 12x DWT histone gene arrays at *ΔHisC^cadillac^* that were confirmed via PCR and nanopore sequencing (Figure 5). Using PCR targeting the junctions created during integration, we can determine whether transgenes are integrated at the distal *attP*, proximal *attP*, or both *attP* sites (Figure 5B,C). We recovered distal, proximal, and double insertions of 4x DWT and 6x DWT transgenes, and a distal insertion of a 12x DWT transgene (Figure 5C). A qualitative assessment of the viability of animals with 4x, 8x, 12x, and 24x copies of DWT histones expressed from integrations into *ΔHisC^cadillac^* is similar to previously described histone gene arrays integrated at VK33, with viable adults eclosing only when histone copy number was 8x or greater (Figure 5C). These data indicate that position effects, at least at *HisC* versus the *attP* site at VK33, are not a significant contributor to expression of transgenic RD histone gene arrays.

**Figure 5.**
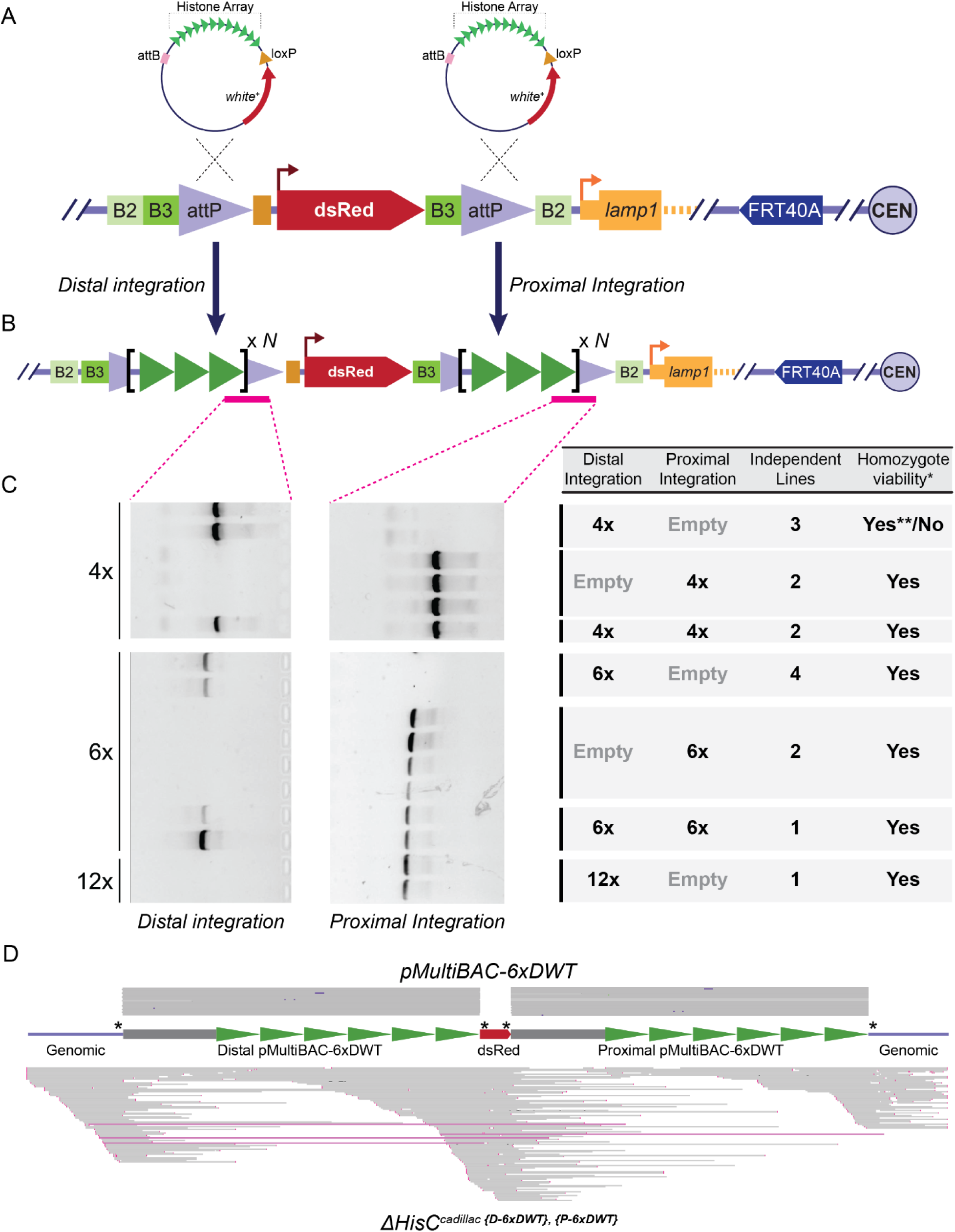
PhiC31-mediated integration of histone gene arrays into *ΔHisC^cadillac^*. A) Diagram of the predicted integration via ΦC31 recombination between the *ΔHisC^cadillac^ attP* sites and the *attB* site on the BAC vector. B) Diagram of resulting locus after integration at both the distal and proximal *attP* sites. PCR target sites spanning the junction between the integrated histone gene array/vector sequence and *ΔHisC^cadillac^* sequence are indicated in with pink bars. C) Representative DNA agarose gels with bands indicating amplification across a vector / *ΔHisC^cadillac^* junction. Summary table (right) describes the proximal and/or distal integration of different histone array multimers, the number of independent lines isolated, and (*) whether the integrated transgene supported viability of *ΔHisC^cadillac {Histone Array}^* homozygotes. “Yes**” indicates that 1 of 3 independent lines is homozygous viable. D) Nanopore validation of a pMultiBAC-6xDWT (top) and the resulting *ΔHisC^cadillac {D-6xDWT}, {P-6xDWT}^* (bottom) transformant. Grey lines indicate aligned long read data from plasmid (top; Plasmidsaurus, Inc. (Eugene, OR)) or transformants (bottom; in house). Both sets of data are aligned to the predicted *ΔHisC^cadillac {D-6xDWT}, {P-6xDWT}^* sequence (middle) including genomic DNA (blue line), pMultiBAC (grey boxes), DWT repeat (green arrows), and dsRed (from *ΔHisC^cadillac^*, red arrowed box). Anchor points for the genomic long read alignment are indicated by asterisks (*) on the cartoon reference genome.

We next sought to determine the identity and structure of the histone gene arrays inserted at *ΔHisC^cadillac^*. However, determining the sequence and copy number of these arrays is challenging due to the lack of accessible, cost-effective, and high-throughput technologies for sequencing large, repetitive DNAs. We have used Sanger sequencing of pMultiBAC plasmids prior to injection to confirm the sequence of engineered histone gene arrays, but this short-read method cannot reliably confirm the sequence of each histone gene unit in a large array of repeated units. Further, the only method currently available to determine the number of histone gene units in a transgenic histone gene array is via southern blotting of genomic DNA, which is expensive, low-throughput, and technically challenging (McKay *et al*. 2015; Zhang *et al*. 2019).

We therefore asked whether leveraging Oxford Nanopore long read sequencing technology could overcome the hurdles associated with these commonly utilized methods to accurately determine the sequence and repeat number of integrated histone gene arrays (Figure 6). We sequenced our 12x pMultiBAC injection constructs using a commercial long read sequencing service and attempted to align raw reads to a reference sequence or perform *de novo* sequence assembly to obtain the sequence of the entire 60kb 12x array. Unfortunately, neither of these established long read sequencing alignment methods was successful. We observed that many reads were misaligned or collapsed into smaller repeats, prohibiting assembly of the entire multi-repeat array (Figure 6A). Given that repetitive histone gene arrays are flanked by unique DNA sequence in the pMultiBAC backbone, we anchored sequencing reads to these unique sequences to allow for proper alignment (Figure 6B). Using this strategy, we could generate consensus sequences without sequence collapses and shifts or obfuscation of mutations. Additionally, as this method uncovers small differences in base calls due to bacterial methylation of the DNA, it provides information to determine whether an apparent mutation is real or the result of methylation (Figure 6C).

**Figure 6.**
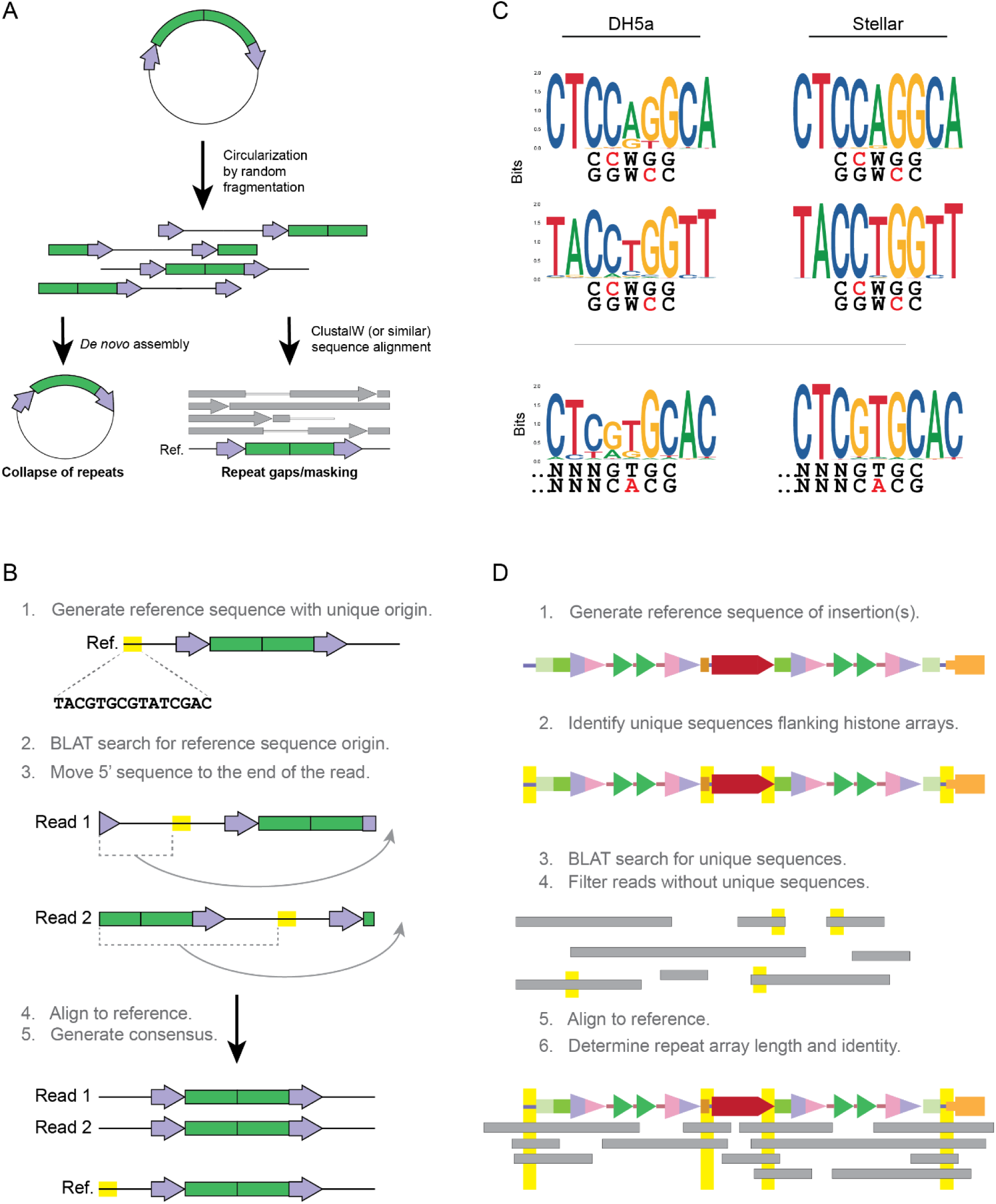
Oxford Nanopore validation of BAC-based histone gene arrays before and after genome integration. A) Commercial plasmid sequencing services rely on the random circularization of plasmids prior to Oxford Nanopore sequencing. *De novo* assembly of these sequences collapse the repeated histone array units (green) while leaving the BAC vector (purple arrows and black line) well annotated. Sequence alignment with ClustalW (or similar) will result in gaps or masking of histone array repeats. B) Schematic of a strategy to determine the histone gene array length and sequence identity using commercial plasmid sequencing with Oxford Nanopore technology. An accurate, linearized reference sequence is generated with repeat regions (green) flanked by unique vector sequence (purple arrows and black line). The first 50-60 bp are selected as the reference origin sequence. A BLAT search is performed against every read, searching for the reference origin sequence. On every read, sequence 5’ of the identified reference origin sequence is moved to the end of the read. The modified reads are aligned, and an accurate reference sequence is generated. C) Oxford Nanopore sequencing will detect prokaryotic DNA methylation as a mis-called base. Logograms showing the depth of base calls around dcm methylation sites (Top) and EcoKI methylation sites (Bottom) grown in either DH5a (dcm+/EcoKI+) or Stellar (dcm-/EcoKI-) *E. coli* cell lines. D) Schematic of a strategy to determine the array length and sequence identity of integrated histone gene arrays using Oxford Nanopore sequencing. An accurate reference sequence of the target locus is generated and ∼50 bp of sequence flanking the integrated histone arrays are selected (yellow highlights). A BLAT search is performed against every read, searching for the unique reference sequences. Reads without these sequences are removed. Alignment with minimap2 generates an accurate map of the sequence.

Having validated this approach using DNA purified from *E. coli*, we next applied it to genomic DNA to determine the structure of the transgenic histone gene arrays integrated in *ΔHisC^cadillac^*. We performed high molecular weight genomic DNA extraction from transformed flies with both distal and proximal 6xDWT integrations in *ΔHisC^cadillac^* (*ΔHisC^cadillac {D-6xDWT}, {P-6xDWT}^*) followed by whole genome sequencing on an Oxford Nanopore GridION. Because reads containing only histone gene array sequence results in ambiguous alignments due to the nature of the repeat, we only aligned reads that overlapped with unique, flanking genomic sequences (Figure 6D, 5D). Using this approach, we identified four reads spanning each 41 kb 6x insertion and >50 reads at each junction between *ΔHisC^cadillac^* and pMultiBAC-6xDWT sequences (Figure 5D). Despite de-enrichment of reads over the histone gene repeats arrays, these data confirm the integration of two complete 6x histone gene arrays in the *ΔHisC^cadillac {D-6xDWT}, {P-6xDWT}^* transgenic line (Figure 5D). Thus, this new approach obviates the need for southern blots to confirm the integration, orientation, and length of transgenic histone gene arrays at *ΔHisC^cadillac^*.

### Tissue-specific expression of mutant histones without using mitotic recombination

Generating mosaic tissue via the mitotic recombination strategy described above offers a way to assess phenotypes of histone mutants in various tissues like the eye. However, clonal assays based on mitotic recombination also have drawbacks, including competition with neighboring wild-type tissue resulting in small populations of mutant cells that are difficult to study. Additionally, generating sister cells with different genotypes via mitotic recombination requires cell division, restricting the analysis of histone mutant cells to growing tissues at certain developmental stages. Moreover, histone mutant phenotypes currently can only be assessed in post-mitotic tissues when the homozygous mutant animal is viable.

To solve these problems, we designed *ΔHisC^cadillac^* to contain recognition sites for the B2 and B3 site-specific recombinases (hereafter B2R and B3R, respectively). These recombinases were originally identified and characterized in yeast and then subsequently engineered for use in *Drosophila* (Toh-e and Utatsu 1985; Utatsu *et al*. 1987; Nern *et al*. 2011; Williams *et al*. 2019). Importantly, B2R and B3R were shown to have robust activity at their specific recognition sites, B2S and B3S, respectively, with no cross reactivity with other recognition sites, including the FLP recognition site, FRT (Nern *et al*. 2011). The specificity of these enzymes allows for the utilization of *B2S*, *B3S*, and *FRT* sites in a single genotype. In *ΔHisC^cadillac^*, the distal *attP* site and *Act5C-dsRed* expression cassette is flanked by *B3S* sites, while the entire *ΔHisC^cadillac^* locus, including *Act5C-dsRed* and both *attP* sites, is flanked by B2S sites (Figure 3E). Thus, expression of B3R will excise the distal *attP* (Figure 7A), while expression of B2R will result in the excision of both *attP* sites (Figure 7B). Each excision event removes *Act5C-dsRed* resulting in loss of *dsRed* expression. To test the efficiency of the B2S/B2R and B3S/B3R in excising *Act5C-dsRed*, we used the UAS/Gal4 system to express either recombinase in third instar wing imaginal discs heterozygous for an empty *ΔHisC^cadillac^*. When expressing B3R in the anterior compartment using ci-Gal4 (marked by GFP+ cells) we found efficient loss of dsRed expression (Figure 7C). Likewise, when expressing B2R with the same driver (marked by GFP+ cells) we also found loss of dsRed (Figure 7D). Together these results indicate that the B2S/B2R and B3S/B2R pairs work efficiently at the *ΔHisC^cadillac^* locus.

**Figure 7.**
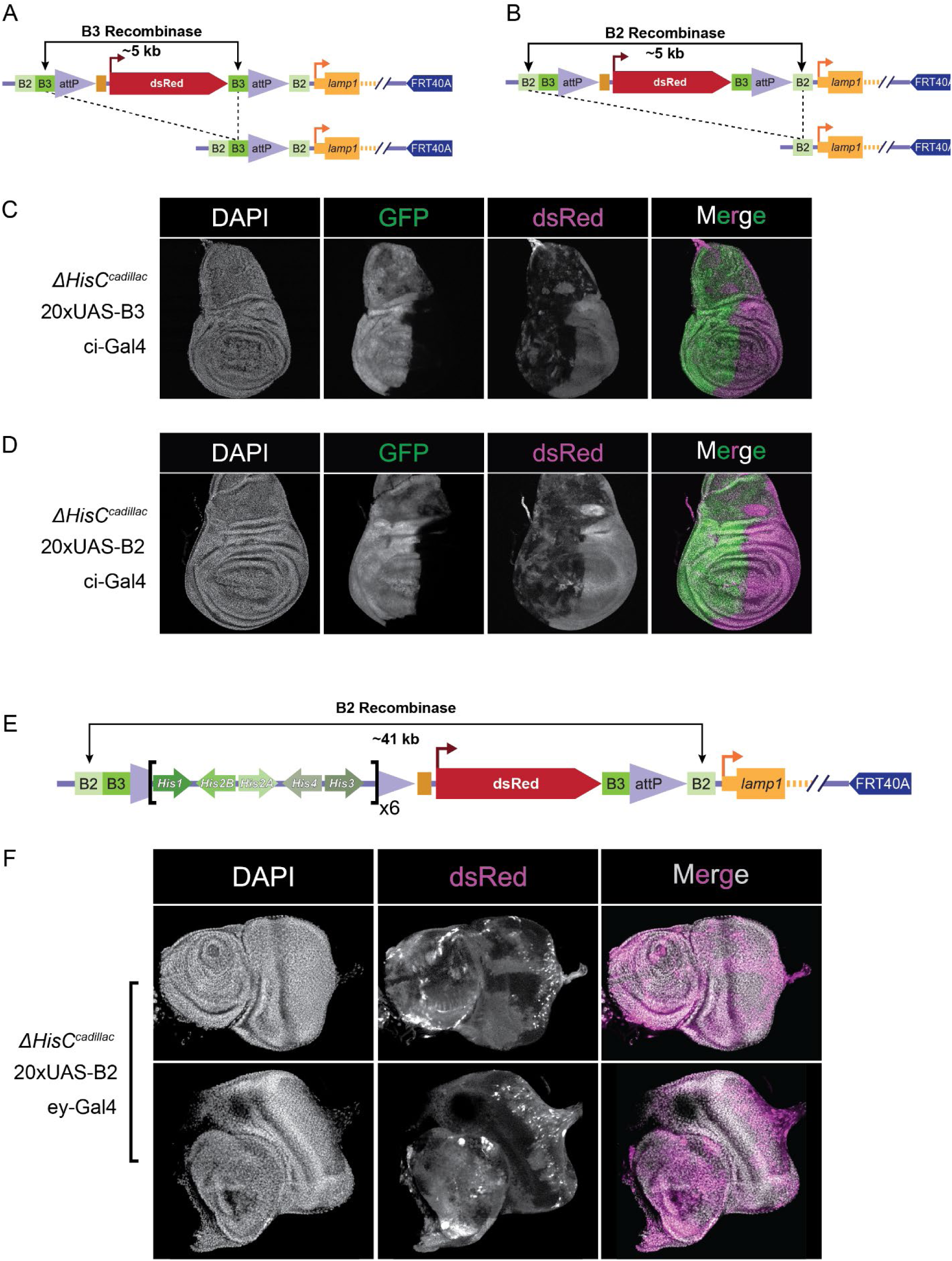
Excision of *ΔHisC^cadillac^* elements and histone gene arrays in a tissue-specific manner. A) Diagram of the predicted 5 kb excision from *ΔHisC^cadillac^* after B3 recombinase expression. B) Diagram of the predicted 5 kb excision from *ΔHisC^cadillac^* after B2 recombinase expression. C) Confocal images of DAPI-stained *Drosophila* third instar wing imaginal discs in which UAS-B3 is expressed in the anterior compartment by Gal4 under the control of the cubitus interruptus (ci) promoter. UAS-sfGFP is also expressed in the anterior compartment and marks cells with recombinase expression. Loss of dsRed signal indicates excision of the ∼5kb dsRed expression cassette from *ΔHisC^cadillac^*. D) As in C except with expression of the B2 recombinase. E) Diagram depicting integration of a 6x wild-type histone gene array into the distal *attP* site of *ΔHisC^cadillac^*, location of the 41kb sequence excised upon expression of B2R is noted. F) Confocal images of two DAPI-stained *Drosophila* eye imaginal discs in which UAS-B2 recombinase is expressed in the full eye disc and part of the attached antennal disc by Gal4 under the control of the eyeless (ey) promoter. Loss of dsRed signal indicates excision of the 41kb dsRed expression cassette plus the 6x wild-type histone array from *_ΔHisCcadillac_*.

We designed *ΔHisC^cadillac^* with the intention of integrating a wild-type histone gene array at the distal *attP* site and a mutant histone gene array at the proximal *attP* site (Supplemental Figure 2A,B). This arrangement of integrated histone gene arrays would allow for the removal of the wild-type histone gene array via B3 excision, leaving intact the mutant histone gene array at the proximal *attP* site and the generation of histone mutant tissue (Supplemental Figure 2C). Note that a similar situation can be generated using mutant histone gene arrays integrated at an ectopic location like VK33 on chromosome 3 (Supplemental Figure 2D). To test this premise, we asked whether a histone gene array integrated in *ΔHisC^cadillac^* could be efficiently excised. We expressed B2R in the eye disc using *ey-Gal4* in an animal with a 6x DWT histone gene array integrated in the distal *attP* site of *ΔHisC^cadillac^* (Figure 7E). Large portions of these eye discs lost dsRed expression, marking the removal of the 6x DWT histone gene array (Figure 7F). However, we found that excision of 6x DWT was not as efficient as removal of the empty 5kb cassette, as clones of dsRed positive cells remained (Figure 7F). Insertion of a 6x wild-type histone array into *ΔHisC^cadillac^* increases the linear genomic distance between the B2S sites by 8-fold (Figure 7E). Previous data suggest that efficiency of FLP recombination decreases as the linear genomic distance between FRT sites increases (Golic and Golic 1996). Thus, the reduced efficiency of 6x DWT excision likely results from the 41kb distance between B2S in this experiment. Although the B2 and B3 recombinase systems are functional in our system, further work is required to fine tune the efficiency of excision of larger insertions to optimize tissue-specific removal of histone gene arrays.

### Recombinase-mediated cassette exchange (RMCE) of histone gene arrays into *ΔHisC^cadillac^*

The *ΔHisC^cadillac^* locus contains two co-linear *attP* sites, offering distinct integration sites for one or two *attB*-containing BACs carrying a histone gene array. These *attP* sites also were designed to be amenable to **R**ecombinase-**M**ediated **C**assette **E**xchange (RMCE), where each genomic *attP* is paired with one of two BAC *attB* sites (Figure 8A) (Schlake and Bode 1994; Bateman *et al*. 2006). This strategy may be desirable in some contexts because it reduces the amount of exogenous DNA at the engineered locus (Figure 8B). To test whether *ΔHisC^cadillac^* was competent for RMCE with histone gene arrays we designed a new vector. Due to the positioning of the *attP* sites relative to *Actin5c-dsRed* in *ΔHisC^cadillac^*, we chose to take advantage of a drug-selection marker to expedite the isolation of transformants. We re-designed the pMultiBAC vector (McKay *et al*. 2015) to include an *hsp40-nptII* gene, a single B3S, PaqCI restriction enzyme sites for Golden Gate cloning (described below), and two co-linear *attB* sites. We named this new BAC-based vector pCadillBAC (Figure 8A). The *hsp40-nptII* confers resistance to food-supplemented G418 sulfate. When a mixed population of larvae consume food containing G418 sulfate, only those expressing *nptII* will develop to adulthood (Matinyan *et al*. 2021a; b). Growing mixed populations of larvae on food with supplemented G418 sulfate without heat-shock was sufficient to select against any flies lacking either a partial (one attP/attB integration, *dsRed+*) or full RMCE (*dsRed-*) of a 4xDWT histone gene array. These data demonstrate that the *ΔHisC^cadillac^* is amenable to RMCE and that drug-based selection methods can be used to supplement or replace visible marker-based methods.

**Figure 8.**
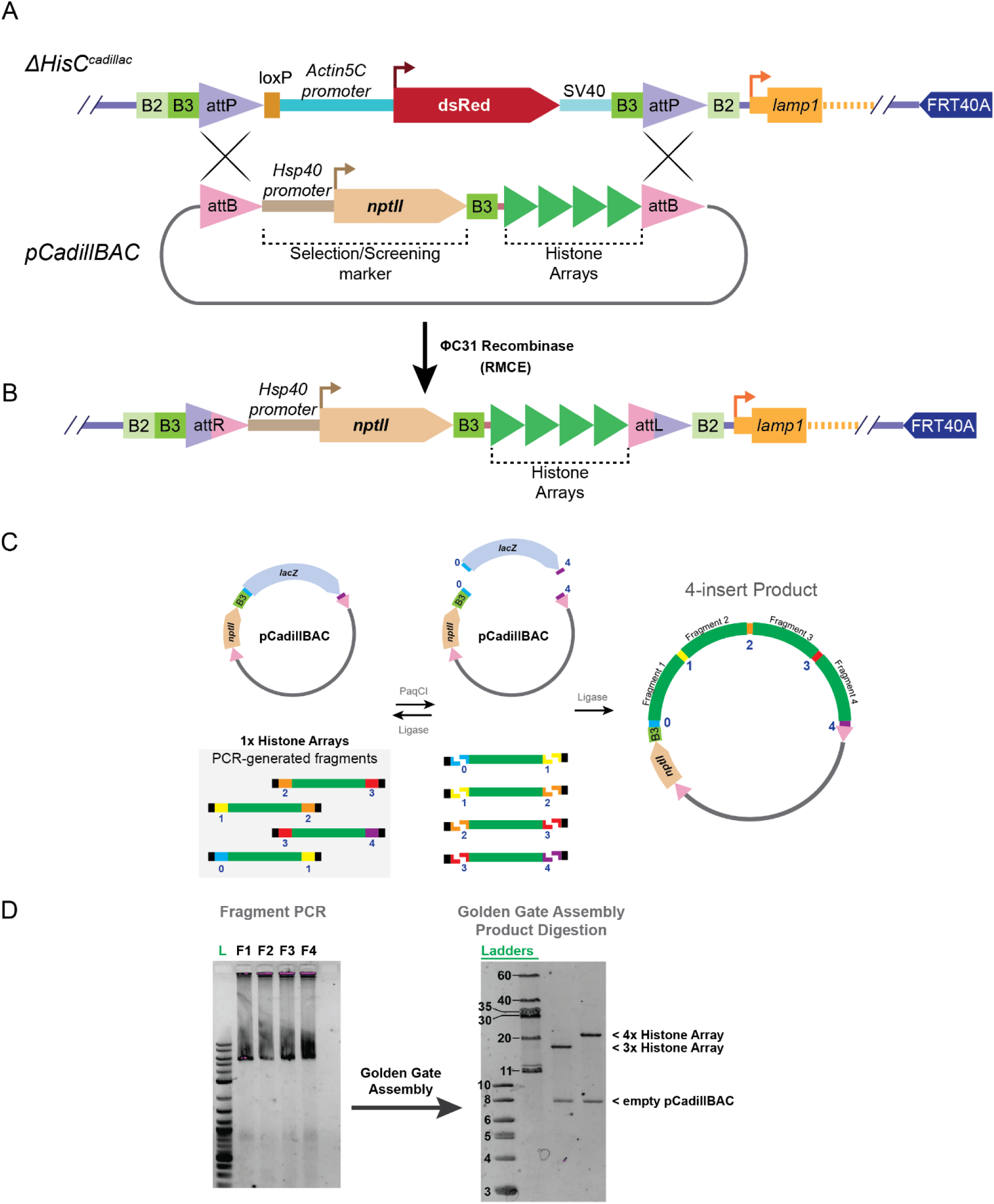
Recombination-mediated cassette exchange (RMCE) of histone gene arrays into *ΔHisC^cadillac^* and rapid generation of histone gene array BACs. A) Diagram of RMCE between pCadillBAC and *ΔHisC^cadillac^*. pCadillBAC is designed with two co-linear *attB* sites, an *hsp40-nptII* selection marker, a B3 site for subsequent selection marker excision, and Golden Gate Assembly-compatible sequences (not shown). B) The resulting locus after RMCE contains little extraneous DNA sequence. C) Diagram of the generation of a 4x histone gene array by Golden Gate assembly through the PCR-generation of 1x histone repeat units (green), reversible cutting of fragments and pCadillBAC, and the final expected product containing the fragments. D) Agarose DNA gel showing generation of each 1x histone gene unit (left) and an agarose DNA gel showing a diagnostic restriction enzyme digest of resultant 3x and 4x histone gene arrays in pCadillBAC.

### Rapid generation of BAC-based histone gene arrays

A bottleneck to generating histone mutant genotypes in all existing systems is the synthesis and sequence confirmation of the large plasmids or BACs containing arrays of histone gene units. We asked whether alternative strategies to traditional restriction enzyme and ligase cloning could be used to generate histone gene arrays. Therefore, we turned to the ‘one-pot’ Golden Gate cloning strategy (Engler *et al*. 2008; Pryor *et al*. 2022a). This strategy utilizes a Type IIS PaqCI restriction enzyme, which generates ends with overhangs of unique sequence because the cut site is distinct from and adjacent to the recognition site. Fragments are designed with compatible ends that when ligated are refractory to re-cutting by PaqCI allowing the assembly and defined arrangement of multiple fragments into single constructs in one reaction (Figure 8D). We designed primers with short indexes to individually PCR-amplify 1x histone gene units (Figure 8E). These PCR reactions generate fragments with compatible ends to either the next array in the sequence or to pCadillBAC. As a result of the PaqCI-ligase reaction, circularized BACs with histone gene arrays are produced (Figure 8D). Via this method, we efficiently produced 4x histone gene arrays in pCadillBAC for RMCE into the *ΔHisC^cadillac^* locus (Figure 8E). Using the pCadillBAC-4xDWT generated with this strategy we successfully recovered transformants that had undergone RMCE at the *ΔHisC^cadillac^* locus. Although we have yet to generate larger histone gene arrays by Golden Gate cloning, likely because of compounded ligation inefficiencies as the number of fragments in the reaction increases, these data provide proof of principle that this rapid cloning strategy can be further developed.

## Discussion

Delineating direct versus indirect phenotypic effects resulting from mutating histone-modifying enzymes and complexes is challenging due to the multitude of substrates targeted by these factors. The approach of mutating histone residues directly is, therefore, a critical capability for more deeply understanding the effects of histone PTMs. *Drosophila melanogaster* is the exemplar animal model organism for studying histone biology with many amenable natural features and engineered tools. Histone replacement platforms were developed by several groups and have been widely used to study different histone residues (Günesdogan *et al*. 2010; McKay *et al*. 2015; Zhang *et al*. 2019; Corcoran and Jacob 2023). Each of these platforms has drawbacks that restrict the types of research questions that can be explored using them. Therefore, we developed a new designer histone locus with expanded capabilities and flexibility. We intend for *ΔHisC^cadillac^* to provide a one-stop-shop for researchers interested in chromatin biology with the flexibility to be customized through integration or modification of the elements.

Our novel histone deletion allele provides several improvements over previous strategies. By using fluorescent markers rather than the *white* gene, we engineered a high-throughput clonal assay in the adult eye to study the effects of histone mutations on cell proliferation and other phenotypes. By modulating H4K16R clone size via mutation of another gene (i.e. *His4r*), we demonstrate that *ΔHisC*^c^*^adillac^* provides a means to employ forward genetics to study histone biology, a first for any animal experimental system. We demonstrated that the *attP* sites in *ΔHisC*^c^*^adillac^* are amenable to proximal, distal, and double integrations of multiple sizes of histone arrays. To reduce the complexity of generating histone mutant arrays and isolating transformants, we developed a new BAC (pCadillBAC) that is compatible with Golden Gate cloning and includes a drug-selectable marker. This BAC is also compatible with RMCE, eliminating extraneous BAC sequence from being added to *ΔHisC^cadillac^*. We also demonstrate site-specific intragenic recombination at *ΔHisC*^c^*^adillac^* using B2 and B3 recombinases. All these functionalities create an opportunity to generate many different histone genotypes in tissue- or temporal-specific ways.

Notably our 6x and 12x DWT integrations represent the first time a transgenic histone array larger than 5x has been integrated into the endogenous *HisC* locus on chromosome 2L. Integration of the transgenic histone gene arrays into *ΔHisC*^c^*^adillac^* is important for mitigating any position effects of ectopic integration sites and to simplify downstream genetic crosses. This system’s full utility will provide the ability to study histone PTM function in different developmental contexts, especially when mutation of histone residues would otherwise result in animal lethality. Furthermore, this system also provides the possibility of investigating histone PTM function in the early embryo and post-mitotic cells through depletion of wild-type histones.

The *ΔHisC*^c^*^adillac^* system is intended to be modular at all levels, allowing for improvements and expansion as new ideas and technologies are developed. The *ΔHisC^cadillac^* locus provides expansion of functionality compared with current histone replacement systems and offers a template for further modulation to fit the needs of individual experiments. We encourage the broader chromatin community to use and modify the *ΔHisC*^c^*^adillac^* system as a resource to further our understanding of epigenetics.

## Data availability

Strains and plasmids are available upon request or in publicly accessible repositories. The authors affirm that all data necessary for confirming the conclusions of the article are present within the article, figures, and tables. Code used to analyze long read data is found at https://doi.org/10.5281/zenodo.11060915 and https://doi.org/10.5281/zenodo.11050973.

## Acknowledgements

We thank Brian Strahl and the UNC histone gene replacement group for fruitful discussions, Jeff Sekelsky for help with Golden Gate cloning and providing cloning vectors, Carolyn Turcotte for help with nanopore sequencing, Susan McMahan for help with expanding the capabilities of *ΔHisC*^c*adillac*^, and Mia Hoover for help with screening for transformants.

## Materials and Methods

### Fly stocks and husbandry

Fly stocks were maintained on standard corn medium provided by Archon Scientific (Durham). The following stocks were obtained from the Bloomington Stock Center:

*y[1] sc[*] v[1] sev[21]; P{y[+t7.7] v[+t1.8]=nanos-Cas9.R}attP2*
*w[1118]; P{y[+t7.7] w[+mC]=20XUAS-B2R.PEST}attP2,*
*w[1118]; P{y[+t7.7] w[+mC]=20XUAS-B3R.PEST}attP2,*
*y[1] w[*]; P{y[+t7.7] w[+mC]=20XUAS-DSCP-B2R}JK65C/TM6C, Sb[1] Tb[1]*
*w[*]; P{w[+m*]=GAL4-ey.H}4-8/CyO*
*y[1] w[67c23]; sna[Sco]/CyO, P{w[+mC]=Crew}DH1*

Other stocks obtained from colleagues:

*y w ; ΔHisC/CyO (*Günesdogan *et al. 2010)*
*y w ; ci-gal4, UAS-GFP (*Uyehara *et al. 2017;* Nystrom *et al. 2020)*

### Cloning

#### ΔHisC^cadillac^ repair template

Homology arms used for CRISPR engineering at *ΔHisC* were PCR-amplified from genomic DNA. The 5’ and 3’ ends of each homology arm are listed below:

##### Distal homology arm (855 bp)

5’genomic sequence – CAACGCGAAAGATTATCATAGATTA
3’genomic sequence - CCCCCTTTTCGGTTGCGCGAGTTGC

##### Proximal homology arm (1193 bp)

5’genomic sequence – GAGACAAAAACGTATGTGTCAGTGT
3’genomic sequence - CGGAAATAAATTGTCCATCAAGTCC

Homology arms were inserted into pBlueSURF (generously provided by Jeff Sekelsky, UNC) by Gibson assembly (NEB). Sequence containing B2 and B3 recombination sites, *attP* sites, *Act5C-dsRed-Express*, and 189 bp of genomic sequence upstream of the *lamp1* transcription start site were synthesized by GENEWIZ, Inc. (South Plainfield, NJ) and cloned between the homology arms in the pBlueSURF construct, generating pBlueSURF-ΔHisC^cadillac^. PAM sites were abrogated using site-directed mutagenesis (NEB). After experimentally determining that the original guide site in the distal homology arm that we tried did not cut efficiently, 59 bp of genomic sequence were added to the 3’ end of the distal homology arm in pBlueSURF-ΔHisC^cadillac^ to allow mutations of the new guide site in the repaired chromosome. Plasmid sequence was confirmed using Sanger sequencing and Oxford Nanopore long read sequencing by Plasmidsaurus, Inc. (Eugene, OR).

#### Guide RNAs used to target ΔHisC

Guide RNA sequences were cloned into pCFD4-U6:1_U6:3tandemgRNAs was a gift from Simon Bullock (Addgene plasmid # 49411; http://n2t.net/addgene:49411 ; RRID:Addgene_49411) (Port *et al*. 2014)

Distal gRNA cloning primer sequence:

5’ - TATATAGGAAAGATATCCGGGTGAACTTCGCAACTCGCGCAACCGAAAAGTTTTAGAGCTAGAA ATAGCAAG - 3’

Distal target sequence:

5’ - GCAACTCGCGCAACCGAAAA - 3’

Proximal gRNA cloning primer sequence:

5’ - ATTTTAACTTGCTATTTCTATTTCTAGCTCAAAACATGCGACGCATTTCATTGCTCGACGTTAAATTGAAAATAGGTC - 3’

Proximal Target sequence:

5’ - AGCAATGAAATGCGTCGCAT - 3’

### CRISPR/Cas9 mutagenesis

Because initial attempts to target *ΔHisC* in the presence of *CyO* failed, likely because of Cas9 cutting at the endogenous histone locus on the *CyO* chromosome, we engineered a stock that expressed Cas9 in the female germline from the *nanos* promoter (P{y[+t7.7] v[+t1.8]=nanos-Cas9.R}attP2) in a *ΔHisC* homozygous background rescued by a homozygous 12x HWT histone gene array. The pBlueSURF-ΔHisC^cadillac^ repair plasmid and pCFD4 containing the proximal and distal gRNAs were injected into *y w*; *ΔHisC*/ *ΔHisC*; {12xHWT}, *nos-Cas9* by Model System Injections (Durham, NC).

We identified transformants that were potential *ΔHisC^cadillac^* repair events by first screening for loss of *white* expression. Of 11 independent transformants that had white eyes, two were also dsRed+, indicating successful integration of the *ΔHisC^cadillac^* sequences. Full sequence of the locus was confirmed by generating a PCR product with the following primers:

5’ - CATGGTATGGGGCGCTATATC - 3’
5’ - GCCGAGAGTCGGTAAAAATG - 3’

followed by TOPO cloning (Invitrogen), and sequencing by Plasmidsaurus, Inc. (Eugene, OR).

### PhiC31-mediated integration of 4x, 6x, and 12x histone transgenes

pMultiBAC containing 4x, 6x, and 12x DWT histone gene arrays were prepared as previously described (Koreski *et al*. 2020), injected into *PhiC31; ΔHisC^cadillac^/(CyO); 8xHWT^w-^/8xHWT^w-^* and screened for *w*+ transformation by Genetivision (Stafford, TX).

The *8xHWT^w-^* chromosome was generated by Cre-mediated excision between the loxP sites in pMultiBAC and the ZH-86Fb landing site, removing pMultiBAC vector sequences, including *mini white* and antibiotic resistance components, and the dsRed marker in 86Fb. Progeny were screened for loss of *white* and dsRed markers.

#### Distal DWT insertions into *ΔHisC^cadillac^* were validated using the following primers

5’ - GTCGCATTTATAAGTGGCAGTGATATATTTTTTGATTGCCAG - 3’
5’ - CTCCGAATATGGCCAGTTGGTC - 3’

#### Proximal DWT insertions into *ΔHisC^cadillac^* were validated using the following primers

5’ - GTCGCATTTATAAGTGGCAGTGATATATTTTTTGATTGCCAG - 3’
5’ - CTAACGAACGTAAGCGACA - 3’

### Generation of w-3xP3-sfGFP and Act5C-sfGFP flies

pattB (generously provided by Jeff Sekelsky) was modified by removing the 5’ region of the *white* gene (including the promoter) and adding either *Act5C-sfGFP* or *3xP3-sfGFP* sequences. *Act5C* promoter sequence was PCR-amplified from pBlueSURF-ΔHisC^cadillac^ plasmid and 3xP3 promoter, sfGFP, and SV40 downstream polyA sequences were PCR-amplified from pScarlessHD-sfGFP-DsRed (Gratz *et al*. 2015) using the following primer sets.

#### Act5C

5’ - GGGAATTGGGAATTCCATGAATGGCATCAACTCTGAATC - 3’
5’ - GCCCTTGGACACCATGGTGGCGTCTCTGGATTAG - 3’

#### 3xP3

5’ - GGGAATTGGGAATTCGGATCTAATTCAATTAGAGACTAATTCAATTAGAGC - 3’
5’ - GCCCTTGGACACCATGGTGGCGACCGGCTTCG - 3’

#### sfGFP

With 3xP3: 5’ - AAGCCGGTCGCCACCATGGTGTCCAAGGGCGAGGAG - 3’
With Act5C: 5’ - TCCAGAGACGCCACCATGGTGTCCAAGGGCGAGGAG - 3’
Shared Rev: 5’ - GAGTCGCGGCCGCTACTTGTACAGCTCATCCATGCCC - 3’

#### SV40

5’ - GAGCTGTACAAGTAGCGGCCGCGACTCTAGATC - 3’
5’ - AGAGGTACCCTCGAGTAAGATACATTGATGAGTTTGGACAAACCAC - 3’

Promoter (Act5C or 3xP3), sfGFP, and SV40 polyA PCR products were assembled in pattB-w-using Gibson assembly (NEB). *Act5C-sfGFP* and *3xP3-sfGFP* integrations into attP40 were recovered by BestGene, Inc. (Chino Hills, CA) and recombined onto an FRT40A-containing chromosome.

### Mosaic eye clones and quantification

*eyFLP; Actin5C-sfGFP/CyO;* + or *eyFLP; Actin5C-sfGFP/CyO; His4r^null^* females were crossed to *yw; ΔHisC^cadillac^/CyO; 12xHTG* or *yw; ΔHisC^cadillac^/CyO; 12xHTG*, *His4r^null^* males. Twenty adults (10 males and females each) were aged at least 1-2 days after eclosion, placed in a 96-well dish containing molten 1% agarose, and cooled to solidify. Images were obtained on a Leica M205 FCA fluorescent microscope using GFP and RFP band pass filters. Quantification of mutant clone size was determined using FIJI (Schindelin *et al*. 2012). Area of the eye from stacked RGB eye images was manually cropped to an ellipse using the selection tool. Mutant eye clones were then white pseudocolored using Color Histogram. The image was then converted into a binary image, followed by restoration of the eye area selection. Finally, the area of the eye covered by mutant clones within the eye area selection was determined.

### Excision from *ΔHisC^cadillac^*

#### Genotypes

**Fig. 7C**: y w; ΔHisC^cadillac^/ci-Gal4, UAS-GFP; P{y[+t7.7] w[+mC]=20XUAS-B2R.PEST}attP2/ +

**Fig. 7D**: y *w; ΔHisC^cadillac^/ci-Gal4, UAS-GFP; P{y[+t7.7] w[+mC]=20XUAS-B3R.PEST}attP2/ +*

**Fig. 7F**: y w; ΔHisC^cadillac{D-6xHWT}^/ey-Gal4; P{y[+t7.7] w[+mC]=20XUAS-DSCP-B2R}JK65C/ +

#### Procedure

Third instar wandering larvae were placed in 1xPBS and mouth hooks were removed with eye discs attached. Mouth hooks and eye discs were fixed in 4% paraformaldehyde (PFA) in 1xPBS for 10 minutes, then washed 3x with 1xPBST (1xPBS with 0.15%TritonX-100). DAPI was added at a concentration of 0.2ug/mL for 10 minutes in 1xPBST, followed by a wash in 1xPBST. Fine dissection of eye discs was performed and all 1xPBST was removed. Discs were mounted on slides with glass coverslips in Prolong Glass (Thermo Fisher Scientific) and imaged using a Leica confocal microscope with a 20X objective. For display in figures all images were separated into channels using FIJI, pseudocolored appropriately, and saved as flat PNGs. Images were then cropped and sized (within groups) to highlight the tissues while maintaining the relative sizing within each figure panel.

### Metazoan histone gene cluster mapping

The approximate locations of the major histone gene clusters in well annotated chromosomes were determined for *H. sapiens, X. laevis, D. rerio,* and *C. elegans.* Briefly, the *Drosophila melanogaster* H2A (FBpp0085249), H2B (FBpp0085281), H3 (FBpp0085250), and H4 (FBpp0085280) protein sequences were collected from Flybase.org (Gelbart *et al*. 1996; Jenkins *et al*. 2022). For each histone protein and each organism, a BLAT search was performed in the UCSC genome browser (hg38, danRer11, xenLae2, ce11; Davis et al., 2022; Homo Sapiens Genome Assembly GRCh38.P14 - NCBI - NLM, n.d.; Howe et al., 2013; Kent et al., 2002; Session et al., 2016) to collect tables of conserved histone proteins and their chromosomal locations. A conservative location of major clusters was identified and chosen for display if more than one histone gene was within 2,500 bp of another. The approximate locations were calculated using the coordinates of the located genes relative to the total length of the chromosomes. The measurements and identification are approximate for global illustration. Cartoon representations of adult metazoans were adapted from BioRender.com

### Metazoan histone protein alignment and conservation

Representative sequences for the canonical histone proteins were collected from UniProt (Bateman *et al*. 2023). The following accession numbers were used: *H. sapiens* H1: P07305, H2A: Q6FI13, H2B: P33778, H3: Q71DI3, H4: P62805; *X. laevis* H1: P22844, H2A: A0A1L8G0Y7, H2B: A0A8J0VAD8, H3: P84233, H4: P62799; *D. rerio* H1: Q6NYV3, H2A: Q561S9, H2B: Q6PC60, H3: Q4QRF4, H4: A3KPR4; *C. elegans* H1: P10771, H2A: Q27485, H2B: Q27484, H3: K7ZUH9, H4: P62784; *D. melanogaster* H1: P02255, H2A: P84051, H2B: P02283, H3: P02299, H4: P84040. Protein sequences were aligned using the ClustalW alignment algorithm in the MegaX program (Thompson *et al*. 1994; Kumar *et al*. 2018). Using the Ident and Sim tool of the Sequence Manipulation Suite (bioinformatics.org/sms2; Stothard, 2000) we determined the percent residue identity and similarity for each species relative to human. Protein alignments were visualized using Alignmentviewer.org (release 1.0) using the EMBL-EBI Mview color scheme (Kanz *et al*. 2005).

### High molecular weight *Drosophila* DNA extraction

Thirty adult flies were collected and frozen and stored at -80°C. Flies were crushed with a tight-fitting pestle in 270 μL Homogenization Buffer (“HB”, 30 mM Tris-HCl pH 8.0, 10 mM EDTA, 0.1 M NaCl, 0.5% Triton X-100). 250 μL additional HB was used to wash the pestle (total volume ∼520 μL). Homogenized flies were centrifuged at 16,000 x g for 5 minutes, before discarding the supernatant. Pellet was resuspended in 520 μL HB by gentle inversion, repeating the centrifugation and resuspension. 60 μL 20% SDS and 20 μL Proteinase K (10 mg/mL) were added followed by incubation at 55°C for 1-4 hours, gently finger flicking every 10 minutes. 1 μL RNase A (100 mg/mL) was added during the final 30 minutes. The mixture was centrifuged at 16,000 xg for 5 minutes and the supernatant carefully transferred into a Phase Lock Gel tube (quantBio 2302830). The supernatant was extracted with 1 volume of phenol:chloroform:isoamyl alcohol twice, and once with equal volume of chloroform:isoamyl alcohol and then the aqueous phase transferred to a 2 mL tube using a wide-bore pipette tip. DNA was precipitated with 0.1 volume of 3M sodium acetate and 2 volumes of ice cold 100% ethanol. DNA was pelleted by centrifugation at 10,000 x g for 5 min. The pellet was washed with 1 mL 70% ethanol twice, before letting it air dry at room temperature for 10 minutes. The pelleted DNA was dissolved in 50 μL nuclease-free water at 4°C overnight, pipetting with a wide-bore 200 μL tip if the pellet remained gelatinous. Qubit and spectrophotometer reading were used to assess DNA quality as outlined in the Oxford Nanopore Native barcoding kit (SQK-NBD114.24).

### Oxford Nanopore long read sequencing of plasmids and genomic DNA

Plasmids were grown in *E. coli* under appropriate conditions and purified using GeneJET plasmid miniprep kit (Thermo Fisher Scientific) using included instructions. Plasmids were verified using Oxford Nanopore long read sequencing services by Plasmidsaurus, Inc. (Eugene, OR) including preliminary annotation by pLann.

Transgenes were verified using Oxford Nanopore long read sequencing performed on the GridION platform. High molecular weight genomic DNA was first extracted (as described above) and the Oxford Nanopore kit SQK-NBD114.24 was used to generate libraries using manufacturer instructions. Libraries were sequenced for up to 72 hours, dependent upon the remaining capacity of the flow cell pores.

To assess the quality of the reads the run report was examined, and reads were assessed using the NanoPlot and NanoStat tools of the NanoPack (De Coster and Rademakers 2023). Reads were also examined for contamination by creating short reads from the 100 bases in the center of each read and running FastQ Screen (Wingett and Andrews 2018). We noted a high amount of *Wolbachia* endosymbiont contamination, with little-to-no contamination from other expected sources (Human, *S. cerevisiae*, *E. coli*). These reads were not expected to our reference sequences, however this suggests that our sequencing runs were hampered by these unnecessary DNA contaminants.

### Generation of accurate Histone gene array plasmid and transgene maps

Alignment visualization and consensus sequence determination required accurate maps of predicted transgenic loci. We provide the sequence of the pCadillBAC vector and *ΔHisC^cadillac^* locus determined by Oxford Nanopore long read sequencing. SnapGene software (www.snapgene.com) was used to assemble histone array plasmids *in silico* using Actions > Restriction and Insertion Cloning or Actions > Golden Gate Assembly > Insert Multiple Fragments. Careful annotation of genes and features was performed manually and by Snapgene > Features > Detect Common Features. These maps are exported as FASTA to be used as reference sequences for later application. An R script (https://github.com/GreshamLab/labtools/blob/master/R/gff_from_snapgene_features.R) was used to convert Snapgene annotations to GFF annotations for later visualization in IGV (Robinson *et al*. 2011).

### Determination of plasmid consensus sequence

Raw FASTQ reads were obtained from Plasmidsaurus, Inc. (Eugene, OR) and run through a custom pipeline to accurately determine the sequence identity and length of histone multimers (https://doi.org/10.5281/zenodo.11050973). Briefly, `seqkit replace` was used to arbitrarily rename reads and `seqkit subseq` was used to select the first 60 bases of a reference sequence (Shen *et al*. 2016). Reads less than 97% and greater than 103% the size of the reference sequence were removed using `seqkit seq`. `seqkit fq2fa` converted the FASTQ to FASTA files. `blat - oneOff=3 -noHead` was used to query the FASTA reads for the beginning of the reference sequence (Kent 2002). `Awk` selected the names of the reads found to contain the reference sequence. Iterating through each read found to contain reference sequence, `seqkit grep` selected the full read and `seqkit restart` reorders the read based on the location of the queried reference sequence. `Seqtk seq` (www.github.com/lh3/seqtk) converted the restarted FASTA reads to FASTQ with arbitrary quality scores. Consensus sequence and alignment was generated using `medaka_consensus` (www.github.com/nanoporetech/medaka) using the restarted reads. Finally, custom Rscripts generate read length histograms, an alignment of the 5’ end of the restarted reads, and bed files that denote homopolymers and bacterial methylation sites which may be incorrectly ascribed to mutations. Resulting aligned reads were visualized on IGV (Robinson *et al*. 2011).

### Determination of integrated transgene sequence

Raw FASTQ reads were obtained by sequencing on an Oxford Nanopore GridION (described above) and run through a custom pipeline to accurately determine the sequence length of transgenic histone multimers (https://doi.org/10.5281/zenodo.11060915). All relevant FASTQ files are concatenated together. Reads are renamed (`seqkit replace`), filtered to a minimum length of 5000 bp (`seqkit seq –min-len`), and converted to FASTA (`seqkit fq2fa`) (Shen et al., 2016). To ensure BLAT does not suffer a segmentation fault, the renamed FASTA reads are split into 20,000 read files with split. Iteratively, each read FASTA file is searched for 50 bp sequences that flank the histone repeat array using `blat - oneOff=3 -noHead`, appending the results to a table (Kent 2002). Matching read names are taken from the table and filtered such that only one instance of each read is kept (`sort names.txt | uniq > blat_names.txt`). Iteratively, `seqkit grep` is used to select each FASTA read that was found to have overlapping sequence. `Seqtk seq` (www.github.com/lh3/seqtk) was used to convert FASTA to FASTQ with arbitrary quality scores. `minimap2 –secondary=no –sam-hit-only -ax -map-ont` was used to map the reads to a reference genome of the transgenic locus (Li 2018). `samtools sort` and `samtools view -bq 1` were used to sort and convert to BAM format and remove multiple-mapped reads (Li *et al*. 2009). Resulting aligned reads were visualized on IGV to find spanning reads (Robinson *et al*. 2011).

### Golden Gate Assembly of Histone Array multimers

Golden Gate Assembly was performed largely according to the manufacturer protocol included with the NEBridge Golden Gate Assembly kit (New England Biolabs) using the with alterations noted below (Potapov *et al*. 2018; Pryor *et al*. 2020, 2022b).

#### Generation of 1x Histone Array fragments

Primers were designed against a 1x DWT histone array (pBS-1xDWT) using SnapGene to include 8 bp unique spacer sequences and PaqCI recognition and cut sites compatible with a 4x multimer in the pCadillBAC vector. The primers below were used with a Q5 High Fidelity Polymerase (NEB #M0491S) PCR to amplify individual histone array amplicons.

##### Fragment 1

5’ - TTGGTCCACCTGCTCCTACCGCTAATGCATATGTGGCGAG - 3’

5’ - TTGGTCCACCTGCTCCTAACAGAGCCGTCTATGTAGTCAAATAAA - 3’

##### Fragment 2

5’ - TTGGTCCACCTGCAAGCTGTTGGCTAATGCATATGTGGCGAG - 3’

5’ - TTGGTCCACCTGCAAGCACTTTAAGCCGTCTATGTAGTCAAATAAA - 3’

##### Fragment 3

5’ - TTGGTCCACCTGCGGTAAAGTCCTAATGCATATGTGGCGAG - 3’

5’ - TTGGTCCACCTGCGGTAAGACGTCGGCCGTCTATGTAGTCAAATAAA - 3’

##### Fragment 4

5’ - TTGGTCCACCTGCTGGTGTCTCTAATGCATATGTGGCGAG - 3’

5’ - TTGGTCCACCTGCATCCCGCCGTCTATGTAGTCAAATAAA - 3’

#### Golden Gate Assembly

Histone array PCR amplicons were incubated with PaqCI (NEB #R0745) for 1-3 hours at 37C and subsequently cleaned up with GeneJet PCR clean up kit (Thermo Scientific) and resuspended in H2O. pCadillBAC vector was incubated with PaqCI for 1-3 hours at 37C, run out on a 1% agarose gel, and gel extracted with GeneJet PCR clean up kit (Thermo Scientific) and resuspended in H_2_O. Equimolar (0.1 pmol) of each digested fragment and vector were mixed and pre-annealed in a thermocycler by heating to 72C and cooling slowly to 15C. Upon reaching 15C, NEBridge Ligase Master Mix (NEB #M1100), NEB PaqCI and PaqCI activator (NEB #R0745) were added and thermocycled per manufacturer recommendations. Assembly mixes were transformed into TransforMax EPI-300 electrocompetent cells (BioSearch Technologies EC300110) and grown on LB + AMP. Correctly assembled vectors were determined by enzyme digest and Oxford Nanopore long read sequencing.

### G418 Selection of transformed progeny

Selection of flies harboring *hsp40-nptII* was performed as described in (Matinyan *et al*. 2021b; a) except that fresh G418 was mixed into the food. Briefly, fly food was gently melted in a microwave and allowed to cool in a double boiler with a stir bar. For every 75 mL of melted fly food, an extra 25 mL of distilled water was added. G418 dissolved in distilled water was added to empty fly vials before adding 20 mL of fly food poured in quickly to mix the solution. Vials were allowed to cool and dry overnight, covered by paper towels. The final concentration of G418 used for selection was 150 mg/mL.

## References

Armstrong R. L., T. J. R. Penke, B. D. Strahl, A. G. Matera, D. J. McKay, et al., 2018 Chromatin conformation and transcriptional activity are permissive regulators of DNA replication initiation in *Drosophila*. Genome Res 28: 1688–1700. 10.1101/gr.239913.118

Armstrong R., T. Penke, S. Chao, G. Gentile, B. Strahl, et al., 2019 H3K9 Promotes Under-Replication of Pericentromeric Heterochromatin in Drosophila Salivary Gland Polytene Chromosomes. Genes (Basel) 10: 93. 10.3390/genes10020093

Baker N. E., K. Li, M. Quiquand, R. Ruggiero, and L.-H. Wang, 2014 Eye development. Methods 68: 252–259. 10.1016/j.ymeth.2014.04.007

Bateman J. R., A. M. Lee, and C. Wu, 2006 Site-Specific Transformation of Drosophila via ϕC31 Integrase-Mediated Cassette Exchange. Genetics 173: 769–777. 10.1534/genetics.106.056945

Bateman A., M. J. Martin, S. Orchard, M. Magrane, S. Ahmad, et al., 2023 UniProt: the Universal Protein Knowledgebase in 2023. Nucleic Acids Res 51: D523–D531. 10.1093/NAR/GKAC1052

Bischof J., R. K. Maeda, M. Hediger, F. Karch, and K. Basler, 2007 An optimized transgenesis system for Drosophila using germ-line-specific φC31 integrases. Proc Natl Acad Sci U S A 104: 3312. 10.1073/PNAS.0611511104

Bongartz P., and S. Schloissnig, 2019 Deep repeat resolution—the assembly of the Drosophila Histone Complex. Nucleic Acids Res 47: e18. 10.1093/NAR/GKY1194

Brown N. P., C. Leroy, and C. Sander, 1998 MView: a web-compatible database search or multiple alignment viewer. Bioinformatics 14: 380–381. 10.1093/BIOINFORMATICS/14.4.380

Chaudhry N., M. Sica, S. Surabhi, D. S. Hernandez, A. Mesquita, et al., 2022 Lamp1 mediates lipid transport, but is dispensable for autophagy in *Drosophila*. Autophagy 18: 2443–2458. 10.1080/15548627.2022.2038999

Copur Ö., A. Gorchakov, K. Finkl, M. I. Kuroda, and J. Müller, 2018 Sex-specific phenotypes of histone H4 point mutants establish dosage compensation as the critical function of H4K16 acetylation in *Drosophila*. Proceedings of the National Academy of Sciences 115: 13336–13341. 10.1073/pnas.1817274115

Corcoran E. T., and Y. Jacob, 2023 Direct assessment of histone function using histone replacement. Trends Biochem Sci 48: 53–70. 10.1016/j.tibs.2022.06.010

Coster W. De, and R. Rademakers, 2023 NanoPack2: population-scale evaluation of long-read sequencing data. Bioinformatics 39. 10.1093/BIOINFORMATICS/BTAD311

Crain A. T., S. Klusza, R. L. Armstrong, P. Santa Rosa, B. R. S. Temple, et al., 2022 Distinct developmental phenotypes result from mutation of Set8/KMT5A and histone H4 lysine 20 in *Drosophila melanogaster*. Genetics. 10.1093/genetics/iyac054

Davis P., M. Zarowiecki, V. Arnaboldi, A. Becerra, S. Cain, et al., 2022 WormBase in 2022—data, processes, and tools for analyzing Caenorhabditis elegans. Genetics 220. 10.1093/GENETICS/IYAC003

Engler C., R. Kandzia, and S. Marillonnet, 2008 A One Pot, One Step, Precision Cloning Method with High Throughput Capability. PLoS One 3: e3647. 10.1371/journal.pone.0003647

Finogenova K., J. Bonnet, S. Poepsel, I. B. Schäfer, K. Finkl, et al., 2020 Structural basis for PRC2 decoding of active histone methylation marks H3K36me2/3. Elife 9. 10.7554/eLife.61964

Gelbart W. M., W. P. Rindone, J. Chillemi, S. Russo, M. Crosby, et al., 1996 FlyBase: The Drosophila database. Nucleic Acids Res 24: 53–56. 10.1093/NAR/24.1.53

Golic K. G., and M. M. Golic, 1996 Engineering the Drosophila Genome: Chromosome Rearrangements by Design. Genetics 144: 1693–1711. 10.1093/genetics/144.4.1693

Gratz S. J., C. D. Rubinstein, M. M. Harrison, J. Wildonger, and K. M. O’Connor-Giles, 2015 CRISPR-Cas9 Genome Editing in *Drosophila*. Curr Protoc Mol Biol 111. 10.1002/0471142727.mb3102s111

Graves H. K., P. Wang, M. Lagarde, Z. Chen, and J. K. Tyler, 2016 Mutations that prevent or mimic persistent post-translational modifications of the histone H3 globular domain cause lethality and growth defects in Drosophila. Epigenetics Chromatin 9: 9. 10.1186/s13072-016-0059-3

Günesdogan U., H. Jäckle, and A. Herzig, 2010 A genetic system to assess *in vivo* the functions of histones and histone modifications in higher eukaryotes. EMBO Rep 11: 772–776. 10.1038/embor.2010.124

Hödl M., and K. Basler, 2012 Transcription in the Absence of Histone H3.2 and H3K4 Methylation. Current Biology 22: 2253–2257. 10.1016/j.cub.2012.10.008

Homo sapiens genome assembly GRCh38.p14 - NCBI - NLM,

Hoskins R. A., J. W. Carlson, K. H. Wan, S. Park, I. Mendez, et al., 2015 The Release 6 reference sequence of the Drosophila melanogaster genome. Genome Res 25: 445–458. 10.1101/GR.185579.114

Howe K., M. D. Clark, C. F. Torroja, J. Torrance, C. Berthelot, et al., 2013 The zebrafish reference genome sequence and its relationship to the human genome. Nature 496: 498–503. 10.1038/NATURE12111

Jacquet K., A. Fradet-Turcotte, N. Avvakumov, J.-P. Lambert, C. Roques, et al., 2016 The TIP60 Complex Regulates Bivalent Chromatin Recognition by 53BP1 through Direct H4K20me Binding and H2AK15 Acetylation. Mol Cell 62: 409–421. 10.1016/j.molcel.2016.03.031

Jenkins V. K., A. Larkin, and J. Thurmond, 2022 Using FlyBase: A Database of Drosophila Genes and Genetics. Methods in Molecular Biology 2540: 1–34. 10.1007/978-1-0716-2541-5_1

Johnston L. A., 2009 Competitive Interactions between Cells: Death, Growth, and Geography. Science (1979) 324: 1679–1682. 10.1126/SCIENCE.1163862/ASSET/04261E34-FBA7-4636-AFF7-AC92240837B6/ASSETS/GRAPHIC/324_1679_F3.JPEG

Kanda H., A. Nguyen, L. Chen, H. Okano, and I. K. Hariharan, 2013 The *Drosophila* Ortholog of *MLL3* and *MLL4* , *trithorax related* , Functions as a Negative Regulator of Tissue Growth. Mol Cell Biol 33: 1702–1710. 10.1128/MCB.01585-12

Kanz C., P. Aldebert, N. Althorpe, W. Baker, A. Baldwin, et al., 2005 The EMBL Nucleotide Sequence Database. Nucleic Acids Res 33: D29. 10.1093/NAR/GKI098

Kaufman T. C., 2017 A Short History and Description of *Drosophila melanogaster* Classical Genetics: Chromosome Aberrations, Forward Genetic Screens, and the Nature of Mutations. Genetics 206: 665–689. 10.1534/genetics.117.199950

Kent W. J., 2002 BLAT--the BLAST-like alignment tool. Genome Res 12: 656–664. 10.1101/GR.229202

Kent W. J., C. W. Sugnet, T. S. Furey, K. M. Roskin, T. H. Pringle, et al., 2002 The Human Genome Browser at UCSC. Genome Res 12: 996–1006. 10.1101/GR.229102

Kimura E. T., M. N. Nikiforova, Z. Zhu, J. A. Knauf, Y. E. Nikiforov, et al., 2003 High prevalence of BRAF mutations in thyroid cancer: genetic evidence for constitutive activation of the RET/PTC-RAS-BRAF signaling pathway in papillary thyroid carcinoma. Cancer Res 63: 1454–7.

Koreski K. P., L. E. Rieder, L. M. McLain, A. Chaubal, W. F. Marzluff, et al., 2020 *Drosophila* histone locus body assembly and function involves multiple interactions. Mol Biol Cell 31: 1525–1537. 10.1091/mbc.E20-03-0176

Kouzarides T., 2007 Chromatin Modifications and Their Function. Cell 128: 693–705. 10.1016/j.cell.2007.02.005

Kumar S., G. Stecher, M. Li, C. Knyaz, and K. Tamura, 2018 MEGA X: Molecular Evolutionary Genetics Analysis across Computing Platforms. Mol Biol Evol 35: 1547–1549. 10.1093/MOLBEV/MSY096

Leatham-Jensen M., C. M. Uyehara, B. D. Strahl, A. G. Matera, R. J. Duronio, et al., 2019 Lysine 27 of replication-independent histone H3.3 is required for Polycomb target gene silencing but not for gene activation. PLoS Genet 15: e1007932. 10.1371/journal.pgen.1007932

Li H., B. Handsaker, A. Wysoker, T. Fennell, J. Ruan, et al., 2009 The Sequence Alignment/Map format and SAMtools. Bioinformatics 25: 2078–2079. 10.1093/BIOINFORMATICS/BTP352

Li H., 2018 Minimap2: pairwise alignment for nucleotide sequences. Bioinformatics 34: 3094–3100. 10.1093/BIOINFORMATICS/BTY191

Lifton R. P., M. L. Goldberg, R. W. Karp, and D. S. Hogness, 1978 The organization of the histone genes in Drosophila melanogaster: functional and evolutionary implications. Cold Spring Harb Symp Quant Biol 42 Pt 2: 1047–1051. 10.1101/SQB.1978.042.01.105

Lindehell H., A. Glotov, E. Dorafshan, Y. B. Schwartz, and J. Larsson, 2021 The role of H3K36 methylation and associated methyltransferases in chromosome-specific gene regulation. Sci Adv 7. 10.1126/sciadv.abh4390

Matinyan N., Y. Gonzalez, H. A. Dierick, and K. J. T. Venken, 2021a Determining effective drug concentrations for selection and counterselection genetics in Drosophila melanogaster. STAR Protoc 2: 100783. 10.1016/j.xpro.2021.100783

Matinyan N., M. S. Karkhanis, Y. Gonzalez, A. Jain, A. Saltzman, et al., 2021b Multiplexed drug-based selection and counterselection genetic manipulations in Drosophila. Cell Rep 36: 109700. 10.1016/j.celrep.2021.109700

McKay D. J., S. Klusza, T. J. R. Penke, M. P. Meers, K. P. Curry, et al., 2015 Interrogating the Function of Metazoan Histones using Engineered Gene Clusters. Dev Cell 32: 373–386. 10.1016/j.devcel.2014.12.025

Meers M. P., T. Henriques, C. A. Lavender, D. J. McKay, B. D. Strahl, et al., 2017 Histone gene replacement reveals a post-transcriptional role for H3K36 in maintaining metazoan transcriptome fidelity. Elife 6. 10.7554/eLife.23249

Meers M. P., M. Leatham-Jensen, T. J. R. Penke, D. J. McKay, R. J. Duronio, et al., 2018a An Animal Model for Genetic Analysis of Multi-Gene Families: Cloning and Transgenesis of Large Tandemly Repeated Histone Gene Clusters, pp. 309–325 in.

Meers M. P., K. Adelman, R. J. Duronio, B. D. Strahl, D. J. McKay, et al., 2018b Transcription start site profiling uncovers divergent transcription and enhancer-associated RNAs in Drosophila melanogaster. BMC Genomics 19: 157. 10.1186/s12864-018-4510-7

Morata G., and P. Ripoll, 1975 Minutes: Mutants of Drosophila autonomously affecting cell division rate. Dev Biol 42: 211–221. 10.1016/0012-1606(75)90330-9

Nern A., B. D. Pfeiffer, K. Svoboda, and G. M. Rubin, 2011 Multiple new site-specific recombinases for use in manipulating animal genomes. Proceedings of the National Academy of Sciences 108: 14198–14203. 10.1073/pnas.1111704108

Nystrom S. L., M. J. Niederhuber, and D. J. McKay, 2020 Expression of E93 provides an instructive cue to control dynamic enhancer activity and chromatin accessibility during development. Development 147. 10.1242/DEV.181909

Pellock B. J., E. Buff, K. White, and I. K. Hariharan, 2007 The Drosophila tumor suppressors Expanded and Merlin differentially regulate cell cycle exit, apoptosis, and Wingless signaling. Dev Biol 304: 102–115. 10.1016/j.ydbio.2006.12.021

Pengelly A. R., Ö. Copur, H. Jäckle, A. Herzig, and J. Müller, 2013 A Histone Mutant Reproduces the Phenotype Caused by Loss of Histone-Modifying Factor Polycomb. Science (1979) 339: 698–699. 10.1126/science.1231382

Pengelly A. R., R. Kalb, K. Finkl, and J. Müller, 2015 Transcriptional repression by PRC1 in the absence of H2A monoubiquitylation. Genes Dev 29: 1487–1492. 10.1101/gad.265439.115

Penke T. J. R., D. J. McKay, B. D. Strahl, A. G. Matera, and R. J. Duronio, 2016 Direct interrogation of the role of H3K9 in metazoan heterochromatin function. Genes Dev 30: 1866–1880. 10.1101/gad.286278.116

Port F., H. M. Chen, T. Lee, and S. L. Bullock, 2014 Optimized CRISPR/Cas tools for efficient germline and somatic genome engineering in Drosophila. Proc Natl Acad Sci U S A 111: E2967–E2976. 10.1073/PNAS.1405500111/SUPPL_FILE/PNAS.201405500SI.PDF

Potapov V., J. L. Ong, R. B. Kucera, B. W. Langhorst, K. Bilotti, et al., 2018 Comprehensive Profiling of Four Base Overhang Ligation Fidelity by T4 DNA Ligase and Application to DNA Assembly. ACS Synth Biol 7: 2665–2674. 10.1021/ACSSYNBIO.8B00333/SUPPL_FILE/SB8B00333_SI_002.ZIP

Pryor J. M., V. Potapov, R. B. Kucera, K. Bilotti, E. J. Cantor, et al., 2020 Enabling one-pot Golden Gate assemblies of unprecedented complexity using data-optimized assembly design. PLoS One 15: e0238592. 10.1371/JOURNAL.PONE.0238592

Pryor J. M., V. Potapov, K. Bilotti, N. Pokhrel, and G. J. S. Lohman, 2022a Rapid 40 kb Genome Construction from 52 Parts through Data-optimized Assembly Design. ACS Synth Biol 11: 2036–2042. 10.1021/acssynbio.1c00525

Pryor J. M., V. Potapov, K. Bilotti, N. Pokhrel, and G. J. S. Lohman, 2022b Rapid 40 kb Genome Construction from 52 Parts through Data-optimized Assembly Design. ACS Synth Biol 11: 2036–2042. 10.1021/ACSSYNBIO.1C00525/SUPPL_FILE/SB1C00525_SI_002.TXT

Regadas I., O. Dahlberg, R. Vaid, O. Ho, S. Belikov, et al., 2021 A unique histone 3 lysine 14 chromatin signature underlies tissue-specific gene regulation. Mol Cell 81: 1766–1780.e10. 10.1016/j.molcel.2021.01.041

Robinson J. T., H. Thorvaldsdóttir, W. Winckler, M. Guttman, E. S. Lander, et al., 2011 Integrative Genomics Viewer. Nat Biotechnol 29: 24. 10.1038/NBT.1754

Salzler H. R., V. Vandadi, B. D. McMichael, J. C. Brown, S. A. Boerma, et al., 2023 Distinct roles for canonical and variant histone H3 lysine-36 in Polycomb silencing. Sci Adv 9. 10.1126/sciadv.adf2451

Schindelin J., I. Arganda-Carreras, E. Frise, V. Kaynig, M. Longair, et al., 2012 Fiji: an open-source platform for biological-image analysis. Nature Methods 2012 9:7 9: 676–682. 10.1038/nmeth.2019

Schlake T., and J. Bode, 1994 Use of Mutated FLP Recognition Target (FRT) Sites for the Exchange of Expression Cassettes at Defined Chromosomal Loci. Biochemistry 33: 12746–12751. 10.1021/bi00209a003

Session A. M., Y. Uno, T. Kwon, J. A. Chapman, A. Toyoda, et al., 2016 Genome evolution in the allotetraploid frog Xenopus laevis. Nature 2016 538:7625 538: 336–343. 10.1038/nature19840

Shen W., S. Le, Y. Li, and F. Hu, 2016 SeqKit: A Cross-Platform and Ultrafast Toolkit for FASTA/Q File Manipulation. PLoS One 11. 10.1371/JOURNAL.PONE.0163962

Shi X., I. Kachirskaia, H. Yamaguchi, L. E. West, H. Wen, et al., 2007 Modulation of p53 function by SET8-mediated methylation at lysine 382. Mol Cell 27: 636–46. 10.1016/j.molcel.2007.07.012

St Johnston D., 2002 The art and design of genetic screens: Drosophila melanogaster. Nat Rev Genet 3: 176–188. 10.1038/nrg751

Stothard P., 2000 The sequence manipulation suite: JavaScript programs for analyzing and formatting protein and DNA sequences. Biotechniques 28. 10.2144/00286IR01

Strahl B. D., and C. D. Allis, 2000 The language of covalent histone modifications. Nature 403: 41–45. 10.1038/47412

Tapon N., K. F. Harvey, D. W. Bell, D. C. R. Wahrer, T. A. Schiripo, et al., 2002 salvador Promotes Both Cell Cycle Exit and Apoptosis in Drosophila and Is Mutated in Human Cancer Cell Lines. Cell 110: 467–478. 10.1016/S0092-8674(02)00824-3

Thompson J. D., D. G. Higgins, and T. J. Gibson, 1994 CLUSTAL W: improving the sensitivity of progressive multiple sequence alignment through sequence weighting, position-specific gap penalties and weight matrix choice. Nucleic Acids Res 22: 4673. 10.1093/NAR/22.22.4673

Tie F., R. Banerjee, P. A. Conrad, P. C. Scacheri, and P. J. Harte, 2012 Histone demethylase UTX and chromatin remodeler BRM bind directly to CBP and modulate acetylation of histone H3 lysine 27. Mol Cell Biol 32: 2323–34. 10.1128/MCB.06392-11

Toh-e A., and I. Utatsu, 1985 Physical and functional structure of a yeast plasmid, pSB3, isolated from *Zygosaccharomyces bisporus*. Nucleic Acids Res 13: 4267–4283. 10.1093/nar/13.12.4267

Tseng A.-S. K., N. Tapon, H. Kanda, S. Cigizoglu, L. Edelmann, et al., 2007 Capicua Regulates Cell Proliferation Downstream of the Receptor Tyrosine Kinase/Ras Signaling Pathway. Current Biology 17: 728–733. 10.1016/j.cub.2007.03.023

Utatsu I., S. Sakamoto, T. Imura, and A. Toh-e, 1987 Yeast plasmids resembling 2 micron DNA: regional similarities and diversities at the molecular level. J Bacteriol 169: 5537–5545. 10.1128/jb.169.12.5537-5545.1987

Uyehara C. M., S. L. Nystrom, M. J. Niederhuber, M. Leatham-Jensen, Y. Ma, et al., 2017 Hormone-dependent control of developmental timing through regulation of chromatin accessibility. Genes Dev 31: 862–875. 10.1101/GAD.298182.117/-/DC1

Weirich S., D. Kusevic, S. Kudithipudi, and A. Jeltsch, 2015 Investigation of the methylation of Numb by the SET8 protein lysine methyltransferase. Sci Rep 5: 13813. 10.1038/srep13813

Williams J. L., H. K. Shearin, and R. S. Stowers, 2019 Conditional Synaptic Vesicle Markers for Drosophila. G3 Genes|Genomes|Genetics 9: 737–748. 10.1534/G3.118.200975

Wingett S. W., and S. Andrews, 2018 FastQ Screen: A tool for multi-genome mapping and quality control. F1000Res 7: 1338. 10.12688/F1000RESEARCH.15931.2

Yung P. Y. K., A. Stuetzer, W. Fischle, A.-M. Martinez, and G. Cavalli, 2015 Histone H3 Serine 28 Is Essential for Efficient Polycomb-Mediated Gene Repression in Drosophila. Cell Rep 11: 1437–1445. 10.1016/j.celrep.2015.04.055

Zhang W., X. Zhang, Z. Xue, Y. Li, Q. Ma, et al., 2019 Probing the Function of Metazoan Histones with a Systematic Library of H3 and H4 Mutants. Dev Cell 48: 406–419.e5. 10.1016/j.devcel.2018.11.047

